# Deep Learning Transforms Phage-Host Interaction Discovery from Metagenomic Data

**DOI:** 10.1101/2025.05.26.656232

**Authors:** Yiyan Yang, Tong Wang, Dan Huang, Xu-Wen Wang, Scott T Weiss, Joshua Korzenik, Yang-Yu Liu

**Affiliations:** Channing Division of Network Medicine, Department of Medicine, Brigham and Women’s Hospital, Harvard Medical School, Boston, MA 02115, USA; Division of Gastroenterology, Hepatology and Endoscopy, Brigham and Women’s Hospital, Harvard Medical School, Boston, MA 02115, USA

## Abstract

Microbial communities are essential for sustaining ecosystem functions in diverse environments, including the human gut. Phages interact dynamically with their prokaryotic hosts and play a crucial role in shaping the structure and function of microbial communities. Previous approaches for inferring phage-host interactions (PHIs) from metagenomic data are constrained by low sensitivity and the inability to accurately capture ecological relationships. To overcome these limitations, we developed PHILM (<u>P</u>hage-<u>H</u>ost Interaction <u>L</u>earning from <u>M</u>etagenomic profiles), a deep learning framework that predicts PHIs directly from the taxonomic profiles of metagenomic data. We validated PHILM on both synthetic datasets generated by ecological models and real-world data, finding that it consistently outperformed the co-abundance-based approach for inferring PHIs. When applied to a large-scale metagenomic dataset comprising 7,016 stool samples from healthy individuals, PHILM identified 90% more genus-level PHIs than the traditional assembly-based approach. In a longitudinal dataset tracking PHI dynamics, PHILM’s latent representations recapitulated microbial succession patterns originally described using taxonomic abundances. Furthermore, we demonstrated that PHILM’s latent representations served as more discriminative features than taxonomic abundance-based features for disease classifications. In summary, PHILM represents a novel computational framework for predicting phage-host interactions from metagenomic data, offering valuable insights for both microbiome science and translational medicine.

## Main

Microbiomes exist in virtually every environment on Earth, from soil and oceans to the surfaces and interiors of plants and animals^1–4^. These communities typically comprise a diverse set of microorganisms, including archaea, bacteria, viruses (notably phages), and fungi, that collectively sustain ecosystem functions or influence host health^5–7^. In the human microbiomes, while numerous studies have documented shifts in prokaryotic taxa between healthy individuals and those with disease^8–13^, the role of phages has become increasingly recognized^14,15^. Emerging evidence indicates that the interactions between phages and their prokaryotic hosts, referred to as phage-host interactions (PHIs) hereafter, are critical determinants in shaping the structure and dynamics of the human microbiomes^16–18^, with profound implications for the development and progression of various diseases^19–24^.

In the absence of comprehensive viral genome catalogs, virome profiling has traditionally relied on *de novo* assembly of shotgun metagenomic reads into contigs for viral identification^25–29^. To infer PHIs, viral contigs are assembled and then phage markers (e.g., clustered regularly interspaced short palindromic repeats (CRISPR) spacers and prophage signatures) are mapped to prokaryotic metagenome-assembled genomes (MAGs)^30,31^. This assembly-based approach of PHI inference has several drawbacks. First, low-abundance phages are under-represented, leading to poor genome coverage and unreliable assemblies^32–34^. Second, PHIs are inferred exclusively from characterized genomic features, restricting predictions to known mechanisms and overlooking novel or uncharacterized relationships. Finally, the high computational demands of the pipelines prolong processing times and increase resource requirements, limiting scalability for large-scale studies.

Recent establishment of human gut virome reference databases^30,35–37^ has driven the development of read-based, reference-dependent virome profilers^32,38^. These tools map viral reads directly to curated catalogs, overcoming the sensitivity limitations of assembly-based approaches and detecting approximately four times more viral species^32^. As a result, they deliver more comprehensive cross-domain profiles of the gut microbiome, accurately representing both prokaryotes and viruses. Moreover, read-based profiling enables unprecedented exploration of PHIs in the human gut through the co-abundance analysis^39,40^. Yet, this co-abundance approach of PHI inference suffers from spurious correlations driven by coincidental co-occurrence rather than genuine biological relationships^41^.

To overcome the limitations of the existing approaches, we investigated deep learning as an alternative approach of inferring PHIs from metagenomic profiles. Deep learning approaches have shown significant potential for elucidating intrinsic, biologically relevant interactions by modeling complex, high-dimensional nonlinear relationships^42–45^. For example, a deep learning model trained on paired microbiome and metabolome datasets accurately predicted metabolomic outputs from microbial compositions and inferred microbe-metabolite interactions^44^. However, no existing deep learning framework has been developed to directly infer PHIs from metagenomic profiles.

In this study, we proposed PHILM (<u>P</u>hage-<u>H</u>ost Interaction <u>L</u>earning from <u>M</u>etagenomic profiles), a deep learning framework that (1) predicts prokaryotic abundance profiles from phage abundance profiles, (2) infers PHIs by performing a sensitivity analysis of the trained deep learning model, and (3) leverages the latent representations learned from the deep learning model as features for downstream tasks (e.g., disease classification and microbial succession pattern tracking) (**Fig. 1**). We tested three deep learning model architectures: NODE (Neural Ordinary Differential Equations), ResNet (Residual Neural Network), and Transformer, and benchmarked them on synthetic data generated from ecological models. We found that all three model architectures outperformed the co-abundance approach in inferring PHIs directly from metagenomic taxonomic profiles. Among them, NODE accurately predicted hosts for 90-97% of phages and recovered more true interactions than both the co-abundance approach and other deep learning model architectures. Therefore, we chose NODE as the representative deep learning model for PHILM in real data analysis. When applied to a dataset over 7,000 healthy-donor stool metagenomes, PHILM uncovered 90% more genus-level PHIs than the assembly-based approach. When applied to a longitudinal dataset tracking PHI dynamics, we further demonstrated that PHILM’s latent representations reproduced the succession patterns originally observed in taxonomic abundances, demonstrating its ability to identify temporal shifts in PHIs. In both real-world datasets, PHILM consistently outperformed the co-abundance approach in recovering known interactions. Finally, PHILM’s latent representations also proved more informative than taxonomic abundance features for distinguishing cases from controls across multiple diseases.

**Fig 1:**
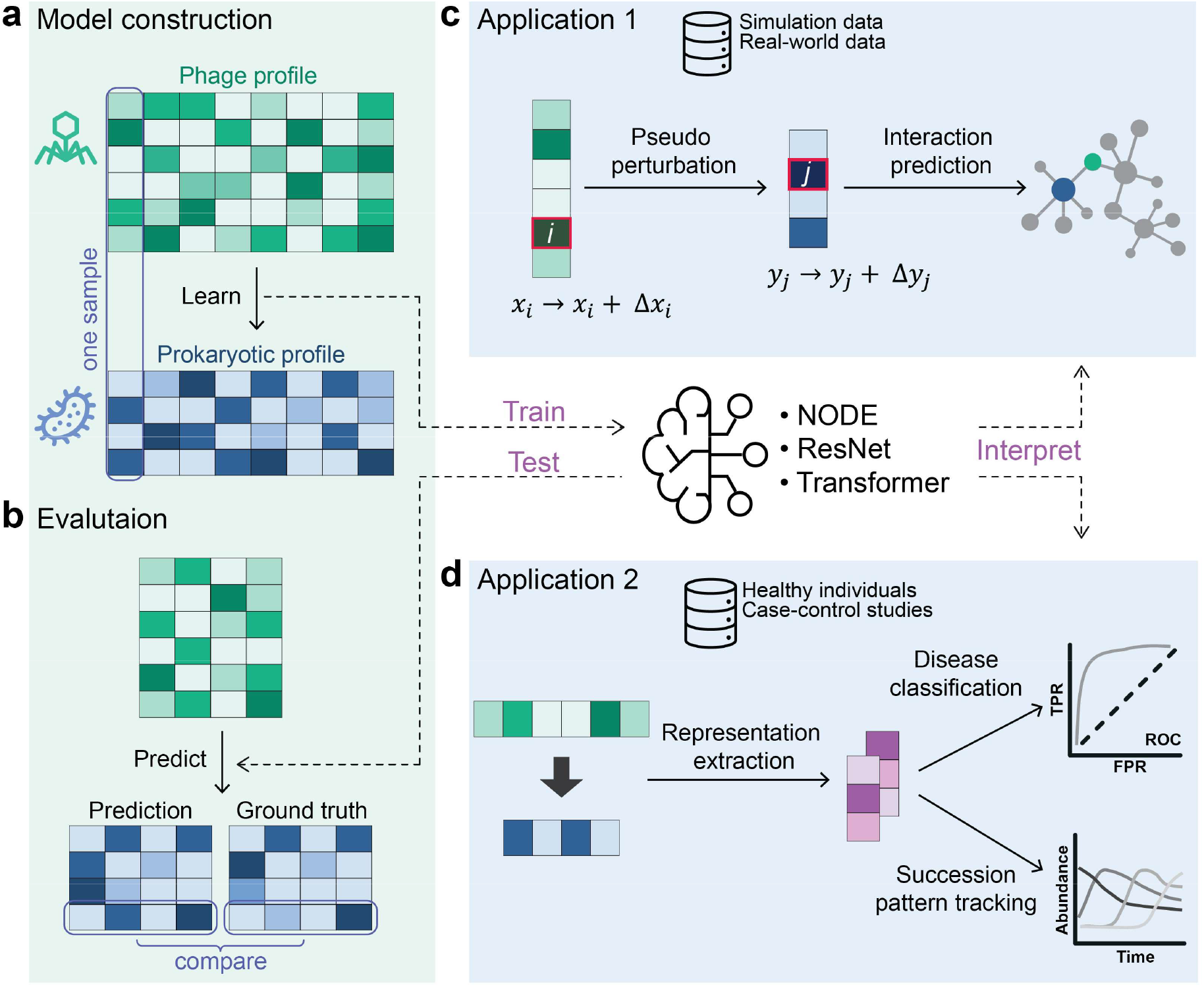
The workflow of the PHILM framework. PHILM is designed to apply deep learning models to discover phage-host interactions from metagenomic data. **a**, Deep learning models are utilized to train on paired phage and prokaryotic relative abundance profiles from the same set of samples. Across all panels, green- and blue-colored matrices represent the phage profiles and prokaryotic profiles, respectively. None of the entities in the matrices has missing data and white cells mean small values. Twelve hypothetical samples with six phage taxa and four prokaryotic taxa are used to illustrate the idea. The samples are divided into training and test sets with a 2:1 ratio. Eight samples in the training set are used for training deep learning models. **b**, A well-trained deep learning model can predict prokaryotic microbial profiles in the test set. For each prokaryotic taxon, the Pearson correlation coefficient (PCC) between its predicted and true abundances across samples in the test set is computed to quantify the models’ performance. **c**, The first application of PHILM is to conduct a sensitivity analysis that perturbs the abundances of input features and predicts alterations in the output. In this way, it helps reveal possible interactions between the input (i.e., phage taxon) and output (i.e., prokaryotic taxon). **d**, The second application involves extracting latent representations from trained models that implicitly encode informative PHIs, which serve as novel features to predict human host phenotypes such as gender in healthy individuals and disease conditions between patients and healthy controls, and to track succession patterns. ResNet: Residual neural network; NODE: Neural ordinary differential equations.

## Results

### Overview of PHILM

For a given set of shotgun metagenomic samples, PHILM takes paired phage and prokaryotic abundance profiles derived from taxonomic profiling as inputs for a deep learning model. In an illustrative dataset of twelve samples, eight were allocated to training and four to testing (**Fig. 1a**). In practice, we randomly split the data into training, validation, and test sets at an 8:1:1 ratio. The validation set serves both for early stopping to prevent overfitting and for hyperparameter tuning to maximize predictive performance. After training, model accuracy is assessed on the test set by computing the Pearson correlation coefficient (PCC) between predicted and experimentally measured relative abundances for each prokaryotic taxon (**Fig. 1b**). The overall predictive power of deep learning models used within PHILM is reported as the mean PCC averaged over all taxa. Previous studies^32,46^ and our analysis of real-world data indicate that phage taxa typically outnumber prokaryotic taxa by a factor of 7∼24 (**Supplementary Table 1**). The high diversity and individual specificity of phages motivated us to utilize phage profiles to predict prokaryotic profiles instead of the other way around.

To evaluate PHILM’s ability to infer PHIs, we conducted a sensitivity analysis (**Fig. 1c, Supplementary Fig. 1a**). We perturbed the relative abundance of phage *i*, denoted as *x*_*i*_, by a small amount Δ*x*_*i*_, and used the trained deep learning model to predict the relative abundance of prokaryote *j*, denoted as *y*_*j*_, and measure the deviation from its original unperturbed abundance, denoted as Δ*Y*. We define the sensitivity of prokaryote *j* to phage *i* as 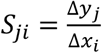. Putative hosts for phage *i* are then identified by one of two criteria: (1) single-host prediction: select the prokaryote *j* with the highest *S*_*ji*_ across samples or (2) multi-host prediction: include any prokaryote *j* for which *S*_*ji*_ exceeds a predefined threshold (**Supplementary Fig. 1b**). More technical details on the computation of *S*_*ji*_ are provided in **Methods**. Furthermore, the latent representations extracted from PHILM can be used to track the microbial succession pattern and serve as more discriminative features than taxonomic abundance-based features for disease classifications (**Fig. 1d**).

### PHILM can predict prokaryotic profiles from phage profiles on synthetic data

To investigate if prokaryotic profiles can be predicted from phage profiles and if PHIs can be inferred from the trained model, we first validated PHILM using synthetic data with known ground truth. The data was generated by two ecological models: the Generalized Lotka-Volterra (GLV) model^47,48^ and the dynamic population (DP) model^49^. Each independent synthetic community (‘sample’) was generated by sampling a set of phages and prokaryotes from a pool of taxa to initiate the community assembly and then collecting the steady-state or dynamically balanced phage and prokaryotic relative abundances as phage and prokaryotic profiles, respectively. Specifically, we assumed a metapopulation pool of *N*_taxa_ = 1,100 (100 prokaryotic taxa and 1000 phage taxa) for both models, with a prokaryote and phage ratio of 1:10 reflecting the taxa ratio we observed in real-world data (**Supplementary Table 1**) and within the estimated ranges from 1:1 to 1:100 reported in previous studies^32,46,50–53^.

The two models differ in complexity and ability to simulate PHIs. The GLV model is relatively simple and ignores the molecular details of phage infection and only accounts for phage-host and inter-host interactions (**Fig. 2a**). We also assumed an extremely narrow-host range for phages during the simulation – each phage taxon can only infect one host, while a bacterial host could be potentially infected by ten phage taxa. Unlike the GLV model, the DP model is more sophisticated and simulates a variety of phage-host processes like infection, lysis/lysogen, induction, reversion and phage death, which is one of the state-of-the-art phage-host dynamic models^54^. In the DP model, each phage is allowed to infect multiple hosts (**Fig. 2b**).

**Fig 2:**
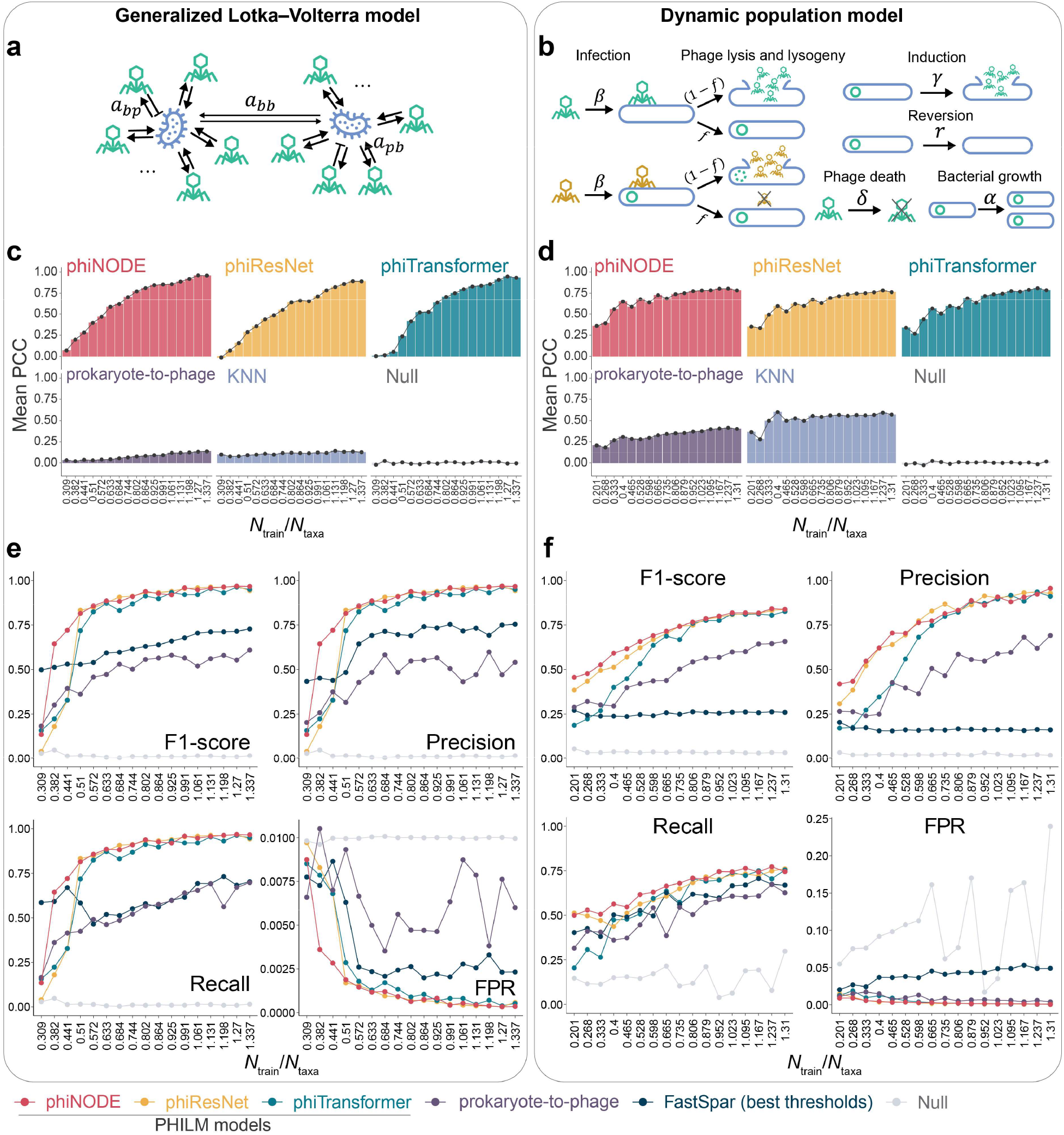
Validation of PHILM using synthetic data. **a**, Schematic of data generation using the generalized Lotka-Volterra model (GLV). *a*_*pb*_ and *a*_*bp*_ represent the phage-prokaryote interactions, while *a*_*bb*_ presents the inter-prokaryotic interactions. Each prokaryote can be infected by 10 phages, and we set strict host specificity for phages, where each phage has only one host. The GLV model was simulated until all phage and prokaryotic abundances reached steady states. **b**, Schematic of data generation using the dynamic population model (DP). Although each prokaryote is set to be targeted by 10 phages, phages are allowed to infect multiple hosts. The DP model was simulated until all phage and prokaryotic abundances reached dynamic balances. Each phage has a lysogen fraction (*f*), indicating its infections leading to lysogeny. Phage with *f* =0 are obligate lytic. More details can be found in the Methods section. **c-d**, Performance of phiNODE, phiResNet, phiTransformer, prokaryote-to-phage method, and a k-nearest neighbors (KNN) model to predict prokaryotic profiles (phage profiles for prokaryote-to-phage method) on the test sets in simulation data from (**c**) GLV and (**d**) DP models with different ratios of training sample size/phage and prokaryote taxa number, *N*_train_/*N*_taxa_. The mean PCCs for all prokaryotic abundances (phage abundances for the prokaryote-to-phage method) across samples are used to reflect the models’ prediction power. **e-f**, Performance of deep learning-based methods and a co-abundance-based method (i.e., FastSpar) to predict PHIs using synthetic data generated by (**e**) GLV and (**f**) DP models with varying *N*_train_/*N*_taxa_ ratios. F1-score, precision, recall, and false positive rate (FPR) were used to evaluate the model’s performance. Note that deep learning methods adopted single-host prediction on GLV model-generated data and multi-host prediction on DP model-generated data. For single-host prediction on DP model data, please refer to **Supplementary Fig. 5**.

We first evaluated the predictive performance of PHILM using three different deep learning architectures (NODE, ResNet, and Transformer) on synthetic datasets generated by the GLV and DP ecological models. The respective models phiNODE, phiResNet, and phiTransformer are named to reflect their underlying model architectures (**Supplementary Fig. 2** and see **Methods**). The mean PCCs for all deep learning models increased with the ratio of training samples to the total number of phage and prokaryotic taxa (*N*_train_/*N*_taxa_). When trained on all available samples with the largest *N*_train_/*N*_taxa_, all deep learning models accurately predicted prokaryotic abundances across samples, achieving mean PCC values of greater than 0.89 on the GLV-generated data (phiNODE: 0.958, phiResNet: 0.890, and phiTransformer: 0.933) and lower values on the DP-generated data (phiNODE: 0.783, phiResNet: 0.764, and phiTransformer: 0.786) (**Fig. 2c-d, Supplementary Table 2**). The relatively low mean Bray-Curtis dissimilarities between predicted and observed community profiles further indicate that these models generally capture prokaryotic composition (**Supplementary Fig. 3**).

We also investigated whether prokaryotic profiles can be used to predict phage profiles by swapping the input and output in the deep learning model (based on NODE and denoted as “prokaryote-to-phage” in **Fig. 2c-d**). When trained with all available samples, the mean PCCs for prokaryote-to-phage are 0.136 for the GLV model and 0.401 for the DP model, which are substantially lower than phage-to-prokaryote predictions. This underperformance likely reflects the challenge of predicting a higher-dimensional output (phage profile) from a relatively lower-dimensional input (prokaryotic profile).

To further confirm the effectiveness of the deep learning models, we implemented a null model using the NODE model architecture but trained on datasets with randomly shuffled sample labels (denoted as “Null” in **Fig. 2**). Across all sample sizes from both GLV and DP models, the null model consistently yielded mean PCCs below 0.035, showing no improvement as *N*_train_/*N*_taxa_ increased. In addition, we tested whether similar phage profiles could predict similar prokaryotic profiles by training k-nearest neighbors (KNN) regressors. For each query phage profile, the model selects its K closest phage profile neighbors and predicts the prokaryotic profile as the distance-weighted average of those neighbors’ prokaryotic profiles. Across varying *N*_train_/*N*_taxa_, the KNN models yield mean PCCs of only 0.08-0.15 for GLV and 0.28-0.59 for DP, indicating that simple inter-profile similarity alone cannot reliably predict prokaryotic profiles (denoted “KNN” in **Fig. 2**). In contrast, the deep learning models consistently achieved higher mean PCCs, demonstrating their effectiveness in predicting prokaryotic profiles from phage profiles, especially when sufficient training samples are available.

Finally, we evaluated whether explicitly modeling inter-prokaryotic interactions could improve prokaryotic profile prediction. Because only the GLV model incorporates inter-prokaryotic interactions, we tested this hypothesis on GLV-generated synthetic data. We appended a graph convolutional network layer to the output of phiNODE, using a known prokaryotic interaction graph to smooth raw predictions before generating the final profiles (see **Methods**). Incorporating these inter-prokaryotic relationships yielded a statistically significant but minor increase in mean PCC (P-value < 0.05; **Supplementary Fig. 4**). However, due to the marginal improvement observed in the synthetic data and the absence of high-confidence inter-prokaryotic interaction networks for empirical datasets, we did not apply this interaction-aware approach to real-world data, although our results demonstrate its potential utility.

### PHILM enables phage-host interaction predictions on synthetic data

Next, we assessed whether PHIs could be inferred from a well-trained PHILM model (e.g., phiNODE, phiResNet, or phiTransformer) by performing a sensitivity analysis (see **Methods**). Notably, we observed that the F1-score, precisions, and recalls of all models increase as the ratio *N*_train_/*N*_taxa_ increases. With the largest *N*_train_/*N*_taxa_, the models accurately reconstruct PHIs, achieving F1-scores no less than 0.94 on the GLV-generated data (phiNODE: 0.966, phiResNet: 0.943, and phiTransformer: 0.949) (**Fig. 2e**). Since the DP model allows phages to infect multiple hosts, we conducted multi-host predictions for all deep learning models by selecting the thresholds for *S*_*ji*_ that maximize the F1-scores across different *N*_train_/*N*_taxa_ ratios. Compared to the performances on the GLV model, lower F1-scores were observed on the DP-generated data (phiNODE: 0.837, phiResNet: 0.837, and phiTransformer: 0.825), reflecting the complexity of the DP model (**Fig. 2f**). With the largest *N*_train_/*N*_taxa_, setting an optimal *S*_*ji*_ threshold of approximately for deep learning model multi-host prediction significantly improved their performance compared to single-host prediction. For example, phiNODE increases its precision from 0.924 to 0.955, recall from 0.670 to 0.745, and F1-score from 0.777 to 0.837 (**Supplementary Fig. 5, Supplementary Table 3**). Again, as expected, the null model achieved an F1-score below 0.06 (**Fig. 2e-f**), which was the lowest among all methods.

We also applied the “prokaryote-to-phage” method with different top phage numbers and optimal thresholds. Although the precisions were moderate, the F1-scores were notably lower (0.61 for the GLV model; 0.66 for the DP model in **Fig. 2e-f** and **Supplementary Fig. 6**). This result demonstrates that predicting phage profiles from prokaryotic profiles is less effective than the phage-to-prokaryote predictions.

In addition to the null model and “prokaryote-to-phage” model, deep learning models were compared with a co-abundance-based model, FastSpar^40^. While FastSpar with a threshold of ±0.5 demonstrated very high precision across different *N*_train_/*N*_taxa_ ratios, it generally exhibited extremely low recall on the GLV-generated data (**Supplementary Fig. 7a**), tending to reconstruct PHIs more accurately for the prevalent prokaryotic species (**Supplementary Fig. 8**). When the threshold was adjusted to ±0.1, FastSpar’s recall improved with small sample sizes but subsequently decreased to below 0.25 as *N*_train_ exceeded 1,200 (**Supplementary Fig. 7a**). However, when evaluated on the DP-generated data, FastSpar with a threshold of ±0.5 exhibited a substantial reduction in precision, with an average of 0.51 (**Supplementary Fig. 7b**). In contrast, FastSpar with a threshold of ±0.1 yielded higher recall but at the expense of precision, averaging only 0.16, and suffered from high false positive rates (**Supplementary Fig. 7b**). It is evident that FastSpar was highly sensitive to the chosen thresholds. Therefore, we selected the thresholds that maximized the F1-scores across different sample sizes for FastSpar. This resulted in modest and slight improvements in F1-scores on GLV and DP model-generated data, respectively (**Fig. 2e-f, Supplementary Fig. 7a-b**). It is worth mentioning that the optimal FastSpar thresholds determined for the largest *N*_train_/*N*_taxa_ were ±0.037 for the GLV model and ±0.048 for the DP model with the largest sample size. These thresholds are substantially lower than FastSpar’s default cutoff of 0.1, potentially leading to high false positive rates, particularly in DP model-based synthetic data (**Supplementary Fig. 7a-b**, the last panels). These findings highlight the limitations of co-abundance-based methods and underscore the superior predictive performance of deep learning models to infer PHIs.

Furthermore, although all the tree PHILM models performed comparably, phiNODE demonstrated superior performance relative to alternative methods in the area under the precision-recall curve (AUPRC) and area under the receiver operating characteristic curve (AUROC), identifying more ground-truth interactions and reaching its peak performance with fewer training samples (**Fig. 2e-f, Supplementary Fig. 9**). On GLV-generated data, phiNODE achieved the best performance, correctly predicting the hosts of 925 out of 958 phages (96.6%). On DP-generated data, it identified hosts for 919 phages (92.4%) using single-host predictions and for 899 phages (90.4%) using multi-host predictions, corresponding to 919 and 1,022 of the 1,371 ground-truth interactions, respectively. Consequently, phiNODE was chosen as the representative PHILM model and used for all subsequent analyses.

### PHILM uncovers 90% more phage-host interactions than the assembly-based approach in microbiome samples from healthy individuals

To evaluate PHILM on real-world data, we downloaded a large-scale dataset comprising 7,016 metagenomic samples of healthy individuals obtained from curatedMetagenomicData^55^ (**Supplementary Table 4**). Taxonomic profiles were generated at the genus level rather than the species level. This decision was informed by our synthetic data analysis, which showed that the F1-score plateaus when *N*_train_/*N*_taxa_ exceeds 1.0 (**Fig. 2e-f, Supplementary Table 3**). Using genus-level taxonomies, we achieved a training sample size (*N*_train_ = 5,612), which is 2.14 times the number of phage and prokaryotic taxa (*N*_taxa_ = 2,620) (**Supplementary Fig. 10a**), and hence is sufficient to ensure reliable interaction prediction. This reliability was further corroborated by a PCC of 0.793 obtained by PHILM on the test dataset (**Supplementary Fig. 10b**). We evaluated PHILM’s sensitivity scores across a range of thresholds and observed that the number of inferred interactions stabilized (relative change < 0.15% over five consecutive thresholds) at a cutoff of 5.0 (**Supplementary Fig. 11**). At this threshold, we recovered 1,906 genus-level PHIs (**Supplementary Table 4**).

To compare with existing methods, we derived PHIs using a conventional assembly-based approach^56^. Specifically, for each sample, contigs were assembled and classified as either viral or non-viral. Non-viral contigs were subsequently binned to generate MAGs of high to medium quality, while viral contigs of medium or higher quality were retained. PHIs were then inferred from genomic features using a strategy similar to that described by Nayfach *et al*. (see **Methods**), which relies on matching prokaryotic CRISPR spacers to viral contigs or performing local alignments between prokaryotic MAGs and viral contigs^56^. This approach yielded over 2,500 MAG-to-contig interactions. To ensure a fair comparison with PHILM, both prokaryotic MAGs and viral contigs were annotated at the genus level, resulting in the identification of 221 genus-level PHIs (**Supplementary Fig. 12**).

In addition to the assembly-based approach, we also applied the co-abundance approach by using FastSpar to infer PHIs with two correlation thresholds: ±0.10 (the default setting) and ±0.09 (the threshold at which interaction counts stabilize) (**Supplementary Fig. 11**). At ±0.10, FastSpar identified 8,111 interactions spanning 285 phage genera; at ±0.09, it detected 9,227 interactions across 334 phage genera, averaging 28 host genera per phage genus. When we compared the overlap of interactions by the assembly-based approach, PHILM, FastSpar (±0.10), and FastSpar (±0.09), we observed 48, 101, and 102 shared interactions, corresponding to positive predictive values (PPVs) of 2.52%, 1.25%, and 1.11%, respectively. Overlapping the PHIs from the Metagenomic Gut Virus (MGV) catalogue^30^ with those from PHILM, FastSpar (±0.10), and FastSpar (±0.09) yielded 239, 363, and 396 common interactions, with PPVs of 12.54%, 4.48%, and 4.29%, respectively (**Fig. 3a, Supplementary Fig. 13a**). To compare PHILM and FastSpar without relying on fixed thresholds, we evaluated both methods across a series of top-N cutoffs. Although they achieved similar overlap with the assembly-based approach, perhaps due to the limited number of interactions detected by this method, PHILM recovered significantly more interactions in common with the MGV catalogue than FastSpar (**Supplementary Fig. 13b-c**). Additionally, specialist-generalist analysis revealed that 71% of phages identified by PHILM are specialists, whereas FastSpar predominantly classifies 73% of phages as generalists with more than three host taxa (**Fig. 3b, Supplementary Fig. 13d**). The much higher proportion of generalist phages predicted by FastSpar may reflect a high false-positive rate, consistent with our benchmarking results using synthetic data (**Fig. 2e-f**). Notably, PHILM identified 89.4% more interactions than the assembly-based approach, potentially suggesting its enhanced sensitivity to dynamic, ecologically relevant phage-host relationships.

**Fig 3:**
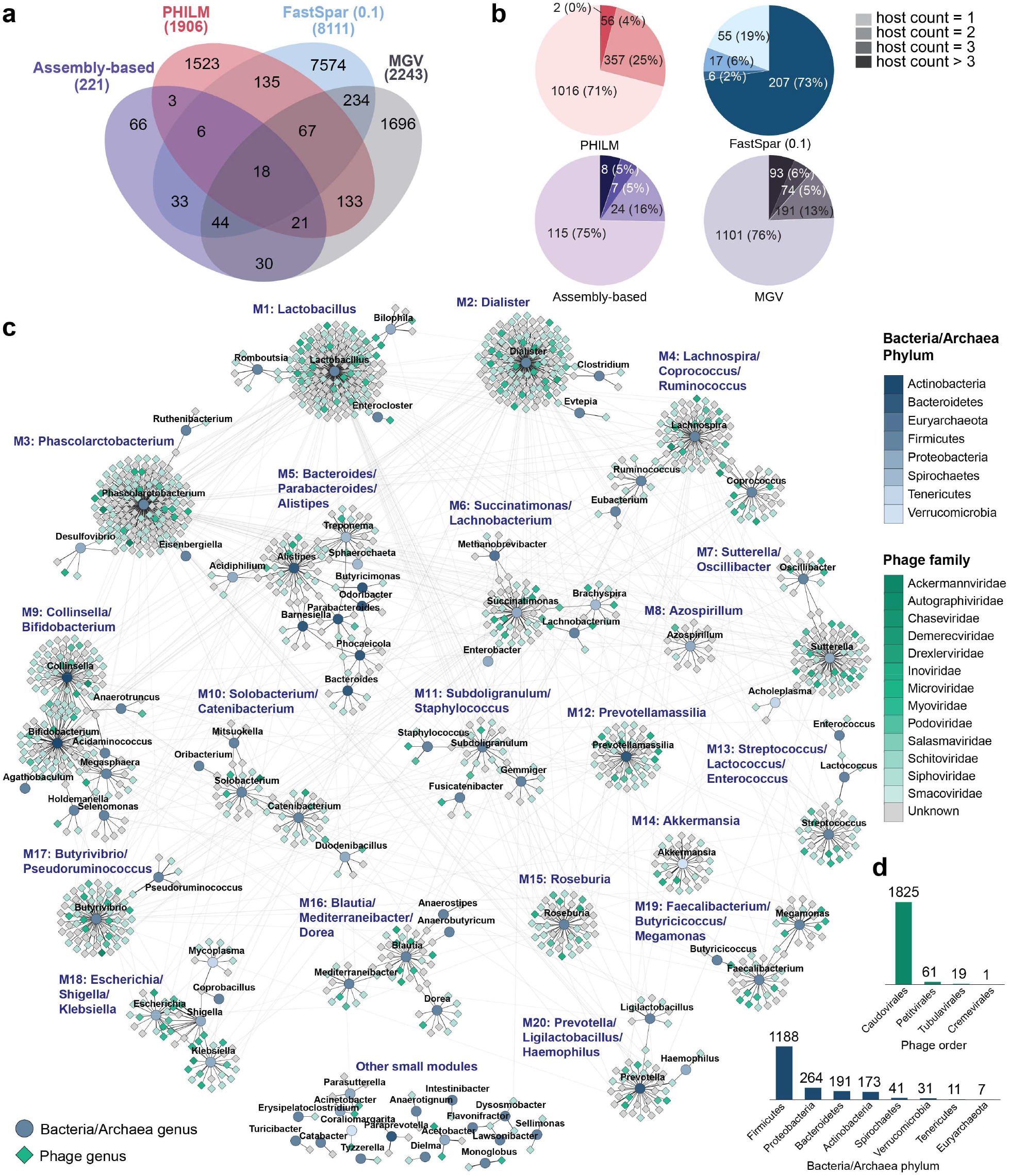
Reconstruction of phage-host interactions in healthy individuals using PHILM. **a**, Venn diagram illustrating the overlaps among PHIs predicted by FastSpar (using a ±0.1 threshold), the assembly-based approach, PHILM, and the interactions recorded in the MGV (Metagenomic Gut Virus) catalogue. **b**, Pie charts illustrating the distribution of phage specificities toward the hosts using different approaches. **c**, PHI networks inferred by PHILM, where circles represent bacteria/archaea genera and diamonds represent phage genera. Phage families and bacteria/archaea phyla are displayed in different colors, and the thickness of the edges indicates the interaction strengths. Nodes that belong to the same modules are clustered together. Edges within modules are shown in black, while those between modules are displayed in light gray. **d**, Distributions of phage orders (top) and bacteria/archaea phyla (bottom) among the interactions identified by PHILM.

Based on PHILM’s predictions, we reconstructed a PHI network and identified 20 densely connected modules (**Fig. 3c**). Several of these modules exhibited clear biological relevance. For instance, module 18 is enriched in pathogenic genera such as *Klebsiella, Escherichia*, and *Shigella*. Module 13 comprises lactic acid-producing cocci, including *Streptococcus, Lactococcus*, and *Enterococcus*. Module 9 contains the age-associated Actinobacteria genera *Bifidobacterium* and *Collinsella* ^57–59^. Module 7 includes *Sutterella* and *Oscillibacter*, both previously linked to obesity^60–63^. Additionally, modules 4, 5, and 20 are dominated by *Ruminococcus, Bacteroides*, and *Prevotella*, respectively, which are genera that define the three gut enterotypes^64^.

The network also revealed that a single prokaryotic host genus can be infected by phages from multiple families, suggesting diverse phages can interact with a relatively narrow host taxonomic range. Furthermore, the prokaryotic hosts in all predicted interactions were predominantly affiliated with the *Firmicutes* phylum, while phages primarily belonged to the *Caudovirales* order, which includes the tailed phage families *Myoviridae, Siphoviridae*, and *Podoviridae* (**Fig. 3d**).

Among the 239 interactions shared by PHILM and the MGV catalogue, only 39 overlapped with those identified by the assembly-based approach (**Fig. 3a**). The remaining 200 exhibited significantly lower, yet predominantly non-zero, abundances in both phage and host genera across samples (**Supplementary Fig. 14**), suggesting that PHILM is more sensitive in detecting low-abundant PHIs. Among these, 154 interactions were missed by FastSpar.

To examine whether genomic features could verify these interactions, we randomly selected seven of the strong PHILM sensitivity values (≥ 9.0). PHILM ranked the *Pseudoruminococcus* genus as the top host for phage genus mgv_g_4001208 despite a weak FastSpar correlation of 0.03 and failed viral contig quality control. Consistently, this association was confirmed by seven *Pseudoruminococcus* genomes in the MGV catalogue and the prophage features found in the host contig (**Fig. 4a**). A similar pattern was observed for mgv_g_4001910-*Treponema* and mgv_g_4000151-*Bifidobacterium*, where PHILM’s high values (10.37 and 9.15, respectively) were supported by MGV records, despite poor assembly-based and FastSpar predictions. The viral and prokaryotic contigs, which were relatively short, aligned well to high-quality phage and host genomes, thereby substantiating these interactions (**Fig. 4b-c**). Interactions mgv_g_4000293-*Sutterella* and mgv_g_4000868-*Megamonas* were not detected by the assembly-based approach due to poor prokaryotic binning but were supported by high PHILM sensitivity values and phage-host genome alignments (**Fig. 4d-e**). In addition, the MGV catalogue lists multiple host genera for these phages—*Sutterella* and *Duodenibacillus* for mgv_g_4000293, and *Megamonas* and *Mitsuokella* for mgv_g_4000868—yet PHILM narrowed and prioritized specific host genera in the healthy human gut. Finally, both PHILM and MGV catalogue uncover the interactions *Brussowvirus*-*Streptococcus* and mgv_g_4001791-*Megamonas*. However, no assembled phage contigs were detected in these cases, highly likely due to extremely low abundances of the phage genera across samples (7.44 × 10^−6^ ± 1.81 × 10^−4^ (mean±standard deviation) for *Brussowvirus* and 2.43 × 10^−6^ ± 6.52 × 10^−5^ for mgv_g_4001791) (**Fig. 4f-g**).

**Fig 4:**
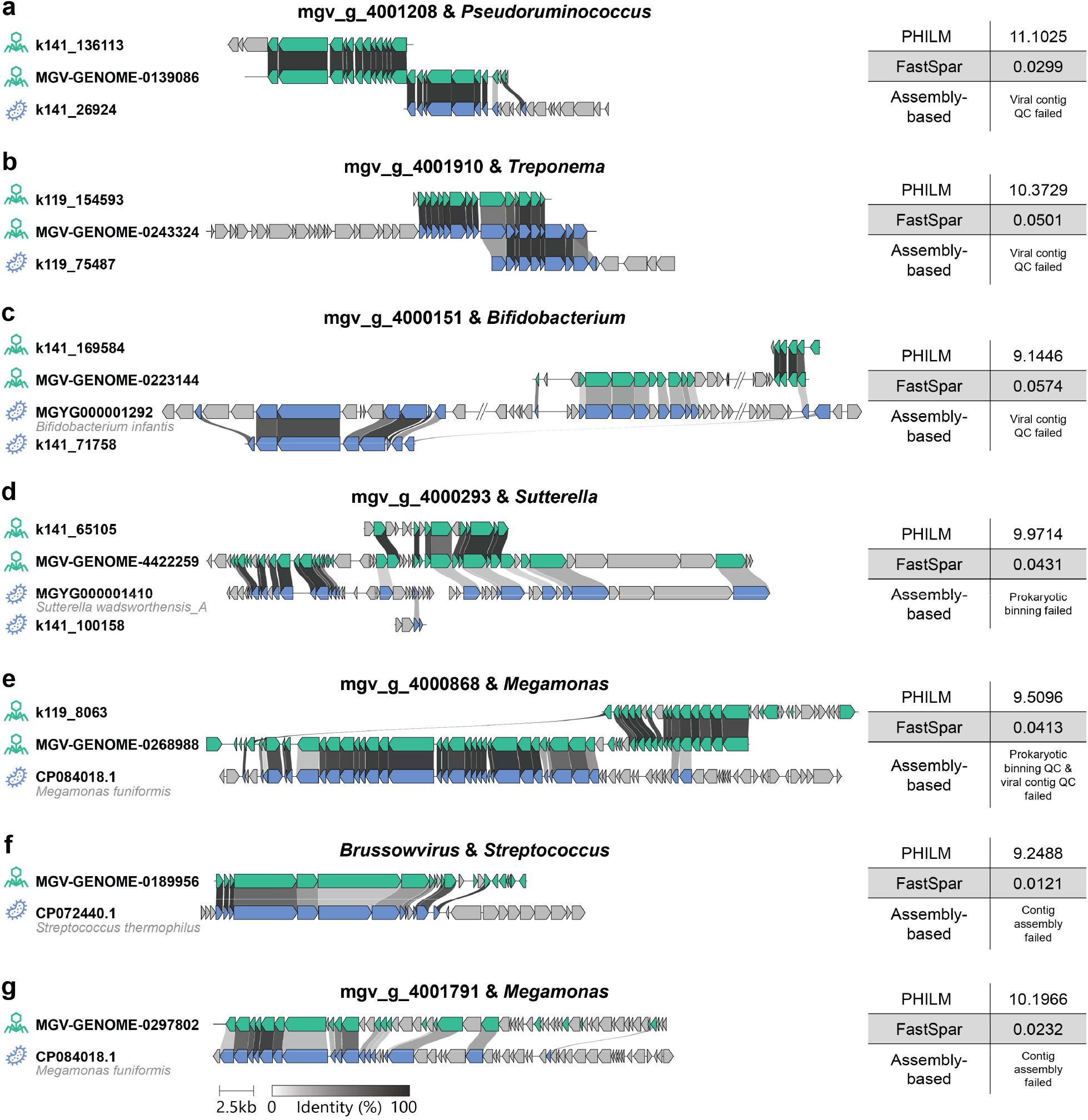
Examples illustrating PHILM’s higher sensitivity in recovering interactions supported by genomic features. **a-c**, Examples show interactions that were not supported by FastSpar or the assembly-based approach, yet are supported by the Metagenomic Gut Virus (MGV) catalogue. **d-e**, Examples illustrate that the interactions were not identified by the assembly-based approach due to poor prokaryotic binning. The MGV catalogue reported multiple prokaryotic host genera for the phage genera, while PHILM prioritized host genera for these phages in the gut of healthy individuals. **f-g**, Examples demonstrate an interaction detected by PHILM in samples that lacked any assembled phage contigs. Although neither FastSpar nor the assembly-based approach identified this interaction, it is supported by the MGV catalogue. The superkingdom icons and genome or contig accessions are shown to the left of each sequence.

Together, these findings demonstrate PHILM’s ability to detect genomically supported, low-abundance, and environment-specific PHIs that both FastSpar and assembly-based approaches overlook.

### PHILM’s latent representations informed by phage-host interactions can be utilized to predict human host phenotypes

Due to insufficient healthy individual stool samples from curatedMetagenomicData for reliably inferring species-level PHIs, which would require over 30,000 samples, we explored alternative applications of PHILM using species-level microbial profiles. Numerous studies have demonstrated the prediction of human host phenotypes using features derived solely from prokaryotic profiles^65,66^. To our knowledge, no prior study has attempted to predict human host phenotypes using phage-inclusive profiles or by incorporating PHIs. PHILM inherently learns PHIs by predicting prokaryotic profiles from phage profiles through the internal layers within its deep learning model (**Fig. 5a**). We hypothesize that these internal latent representations can serve as informative features for predicting human host phenotypes.

**Fig 5:**
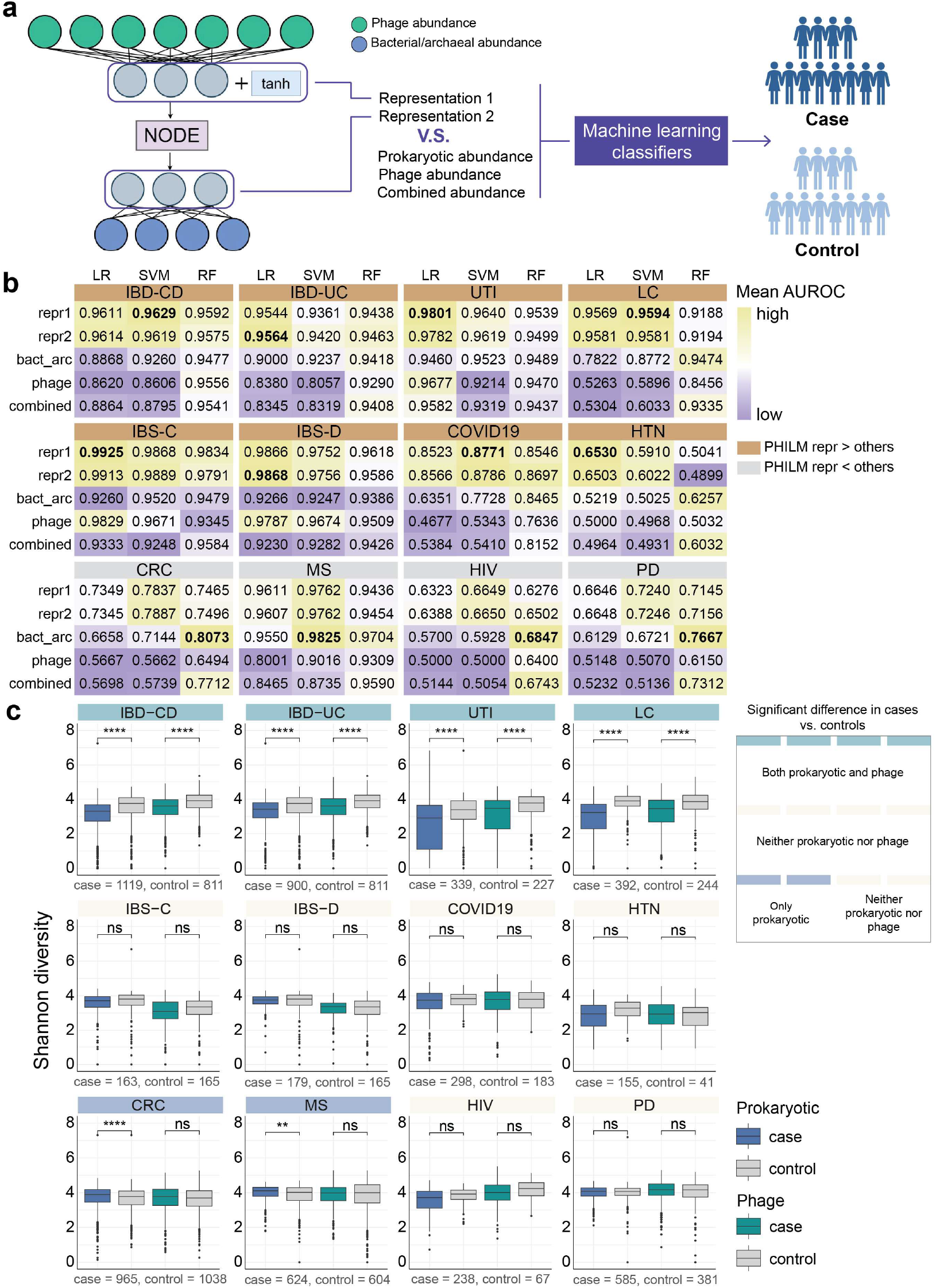
Application of PHILM’s latent representations for case-control classification across diseases. **a**, For a trained PHILM deep learning model (phiNODE shown as an exemplar model) to predict disease-related prokaryotic profiles from phage profiles, its hidden layers implicitly reflecting PHIs can be regarded as latent representations, which can be fed into machine learning-based classifiers as features to distinguish healthy individuals and diseased individuals. Representation 1 (repr1) is the representation close to the input layer (phage profile), while representation 2 (repr2) is close to the output layer (prokaryotic profile). **b**, Heatmap illustrating the mean areas under the receiver operating characteristic curve (AUROCs) from five-fold cross-validation of three machine learning classifiers using PHILM-derived representations, prokaryotic profiles, phage profiles, and combined cross-domain profiles to classify disease conditions across 12 diseases. Higher AUROC values are shown in yellow and lower values in purple, with color scales determined individually for each disease. Green headers denote that PHILM-derived representations achieve the best performance, whereas grey headers indicate that other features perform the best. Rows represent five feature types across different diseases, and columns represent Logistic Regression (LR), Support Vector Machine (SVM), and Random Forest (RF) classifiers. **c**, Boxplots illustrating Shannon diversity indices based on prokaryotic (blue and gray boxes) and phage (green and gray boxes) profiles, comparing cases and controls across various diseases. Sample sizes for cases and controls for each disease are indicated at the bottom of each panel. We employed two-sided Wilcoxon rank-sum tests to assess differences between groups. Significance levels are denoted as follows: ns (p > 0.01), ^**^ (p≤0.01), ^***^ (p≤0.001), and ^****^ (p≤0.0001). IBD-CD: Crohn’s disease; IBD-UC: Ulcerative colitis; UTI: Urinary tract infection; LC: Liver cirrhosis; IBS-C: Irritable bowel syndrome with constipation; IBS-D: Irritable bowel syndrome with diarrhea; HTN: Hypertension; CRC: Colorectal cancer; MS: Multiple sclerosis; PD: Parkinson’s disease.

Given that human host gender and age metadata were available for nearly all stool samples (98.8% for gender and 100% for age in **Supplementary Fig. 10e-f, Supplementary Table 5**), we applied the PHILM-derived latent representations along with other microbial abundance features to three classifiers (Random Forest (RF); Logistic Regression (LR); Support Vector Machine (SVM)) to predict different gender and age groups. Notably, significant differences in Shannon diversities for both prokaryotic and phage compositions were observed between different gender and age groups (**Supplementary Fig. 10g-h**), suggesting that phages may also contribute to distinguishing between these groups. Two representations were extracted from the PHILM’s trained phiNODE model: one from a hidden layer closer to the input (representation 1, repr1) and one from a hidden layer closer to the output (representation 2, repr2) (**Fig. 5a**).

Among all model-feature combinations, the SVM with PHILM repr2 and repr1 achieved the highest average AUROC of 0.943 and the highest mean F1-score of 0.886, respectively, using five-fold cross validation (**Supplementary Fig. 15a-b**). Regarding age prediction, where the data were categorized into five qualitative groups based on established criteria^65^, the SVM with PHILM repr1 and repr2 achieved the best average AUROC of 0.950 and the best mean F1-score of 0.777, respectively, across all strategies (**Supplementary Fig. 15c-d**). Collectively, these results demonstrate that PHILM-derived latent representations exhibit higher predictive power for human host phenotypes than other types of features.

### PHILM recapitulates phage-host dynamics in the early-life gut microbiome

We further evaluated PHILM on a dataset of 12,262 longitudinal stool samples collected from 887 children during early life^46^. In the original study, genus-level PHIs were predicted by iPHoP^31^ by pairing shotgun read-mapped viral and bacterial species-level genome bins from a reference database. Because applying an assembly-based approach to such a large dataset is computationally prohibitive, we compared PHILM only with the co-abundance-based approach (e.g., FastSpar). For both PHILM and FastSpar, we ranked their inferred interaction by strength and measured their overlap with the iPHoP predicted interactions across a series of top-N cutoffs. At every threshold tested, PHILM recovered significantly more iPHoP-validated interactions than FastSpar (**Supplementary Fig. 16**), demonstrating stronger concordance with genome-based inference.

Next, we examined the latent representations learned by PHILM, trained on species-level taxonomic profiles and presumed to encode underlying phage-host interplays, for their ability to capture temporal dynamics. Remarkably, when clustering these embeddings along the day of life, we observed eight distinct subsets whose trajectories (**Fig. 6b-c**) closely aligned with microbial succession patterns reported in the original study (**Fig. 6a**). These findings highlight that PHILM not only infers static PHI networks but also learns representations that capture the temporal dynamics of gut microbiome development.

**Fig 6:**
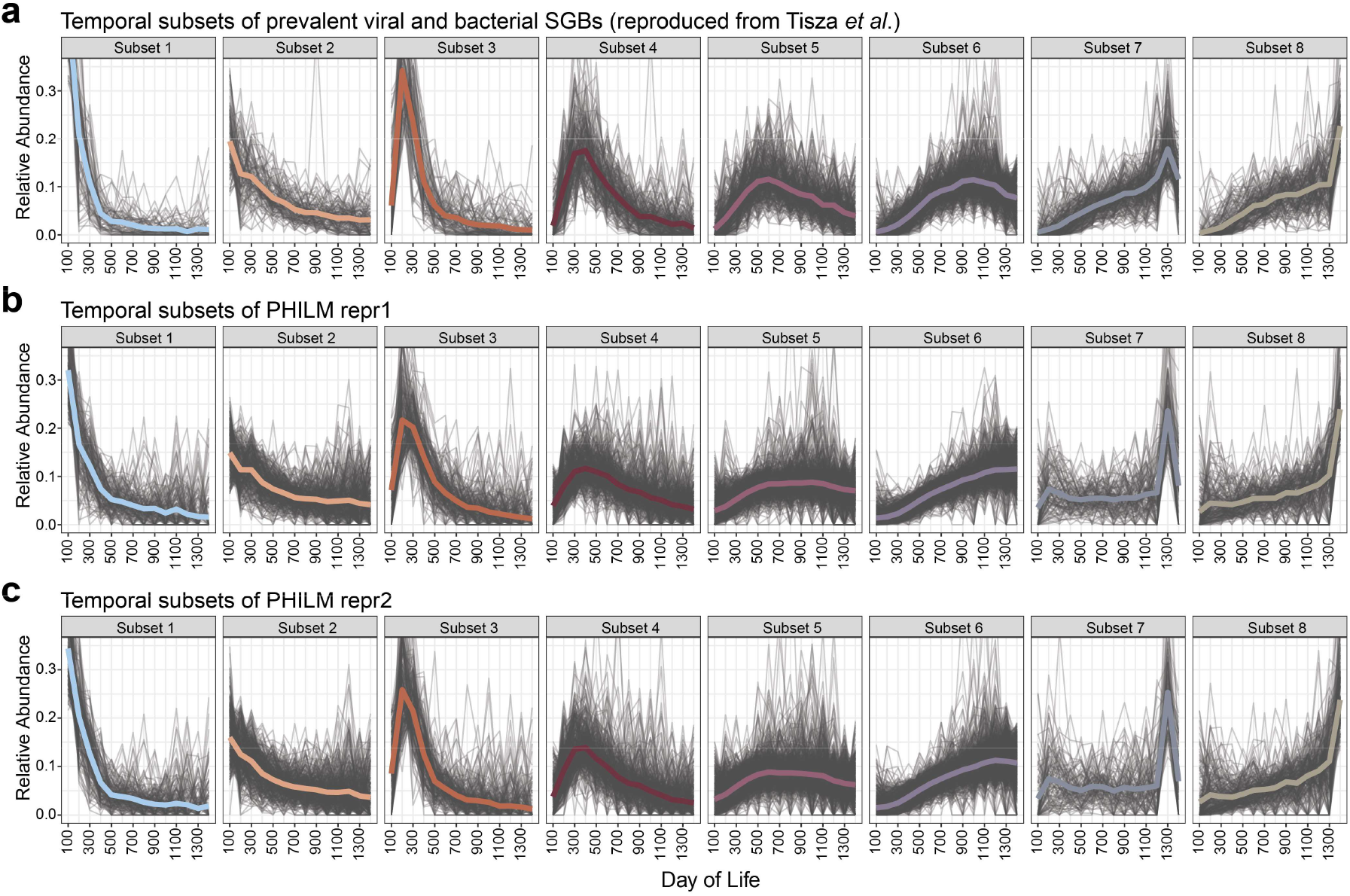
Application of PHILM’s latent representations to reconstruct phage-host dynamics in the early-life gut microbiome. **a**, Temporal subsets of prevalent viral and bacterial species-level genome bins (SGBs), reproduced from Tisza *et al*. Each trajectory shows the abundance changes of a single SGB over days of life. Gray lines represent individual SGBs, and colored lines indicate their average trends. Viral and bacterial SGBs are plotted together, rather than separately in the original study. **b-c**, Temporal subsets of PHILM’s latent representations 1 and 2, respectively. Values in representation vectors were inverse-centered log-ratio-transformed into abundance-like values to facilitate comparison with the original raw abundances.

### PHILM’s latent representations serve as promising features for disease classification

Building on the promising results of using PHILM-derived latent representations for phenotype prediction in healthy individuals, we next evaluated whether these representations could also enhance disease classification. While numerous attempts have been made to differentiate cases from controls using microbial features, most studies consider these features in isolation without incorporating cross-domain data or interactions, specifically phage-host relationships^67,68^. To demonstrate PHILM’s potential in clinical diagnosis, we collected ten shotgun metagenomic datasets representing twelve diseases, including inflammatory bowel disease (IBD), urinary tract infection (UTI), liver cirrhosis (LC), irritable bowel syndrome (IBS), COVID-19 infection, hypertension (HTN), colorectal cancer (CRC), multiple sclerosis (MS), human immunodeficiency virus (HIV) infection and Parkinson’s disease (PD). Cross-domain taxonomic profiles at the species level for both prokaryotes and phages were generated for all datasets (**Supplementary Fig. 17a**), which were used to train PHILM and extract two latent representations (**Fig. 5a**).

We first assessed differences between cases and controls across diseases using permutational multivariate analysis of variance (PERMANOVA). In PERMANOVA, the F statistic represents the ratio of the variance between groups to the variance within groups, with higher values indicating greater inter-group variance than that of intra-groups. We found that the F values for PHILM representations (either repr1 or repr2) in 7 out of 10 datasets are higher than those obtained using prokaryotic features, phage features, or combined features (**Supplementary Fig. 18**). Similar to the previous analysis conducted for the healthy individual dataset, we next used these features as input variables for three classifiers (RF, LR, and SVM) to predict disease conditions. Compared with alternative features, either repr1 or repr2 achieved the highest average AUROCs from five-fold cross-validation in 8 out of 12 diseases (**Fig. 5b, Supplementary Fig. 19**), and they obtained the highest average F1-scores in 7 diseases (**Supplementary Fig. 20, Supplementary Table 6**). Among all models, LR most consistently achieved the best AUROCs. We also noticed that the diseases for which PHILM-derived representations outperformed other features in PERMANOVA did not always align with those showing superior performance in machine learning prediction, suggesting that PHILM representations may better capture non-linear and subtle patterns than traditional statistical tests.

We next investigated why PHILM-derived representations underperformed in four diseases: CRC, MS,HIV infection, and PD. We examined potential factors and found that it could not be attributed to case-control imbalance. Actually, except for HIV, COVID-19, and HTN, most datasets maintained balanced case-control ratios (**Supplementary Fig. 17b**). Additionally, differences in model fitting are unlikely to be the cause, as the mean PCCs for the four underperforming diseases were comparable to those for the others; notably, MS had a mean PCC of 0.73, the second highest among all diseases (**Supplementary Fig. 17c**). We then examined whether alpha diversity differs between cases and controls when using prokaryotic and phage profiles. For diseases showing significant differences in both profiles, such as IBD-CD, IBD-UC, UTI, and LC, PHILM-derived representations achieved the highest mean AUROCs among all features (**Fig. 5c**, the first row). For diseases without significant differences between cases and controls using any profiles, PHILM representations still outperformed other features in four diseases (IBS-C, IBS-D, COVID-19, and HTN) (**Fig. 5c**, the second row), whereas prokaryotic abundance-based features performed the best in the remaining diseases, HIV and PD (**Fig. 5c**, right two panels on the third row). Another scenario where PHILM underperformed was when significant differences were found only in prokaryotic profiles but not in phage profiles (**Fig. 5c**, left two panels on the third row). In other words, PHILM-derived representations consistently outperform other features in disease classification when both prokaryotic and phage profile Shannon diversities differ significantly between cases and controls.

Notably, PHILM’s latent representations reduce dimensionality by roughly two orders of magnitude compared to phage or combined abundance-based features. They were the smallest in seven datasets and the second smallest in the remaining three (**Supplementary Fig. 17d**). This substantial reduction in feature numbers, especially compared to combined abundance features, highlights a more efficient approach to integrating cross-domain species data. Overall, our findings demonstrate that latent representations learned from PHILM outperform traditional species-based features in predicting most disease states, underscoring their potential as robust, low-dimensional biomarkers for future research.

## Discussion

Existing assembly-based approaches for inferring PHIs from shotgun metagenomic data tend to miss many interactions because assembled genomes poorly represent low-abundance phage taxa^32–34^. Moreover, these methods rely heavily on known molecular mechanisms, such as infection processes or host defense systems, which limit their ability to discover novel PHIs. In this study, we developed PHILM, a deep learning framework that predicts prokaryotic profiles from paired phage profiles. By interpreting the trained model, we infer PHIs in a purely data-driven fashion without relying on any predefined biological mechanisms, potentially capturing more phage-host ecological dynamics. When tested on synthetic data, PHILM outperformed co-abundance-based methods, while applied to a large, cross-cohort collection of >7,000 healthy individual stool samples, PHILM identified more plausible PHIs and a higher proportion of specialist phages than the co-abundance approach. More importantly, it recovered substantially more interactions than the traditional assembly-based pipeline, which enabled a biologically meaningful reconstruction and modulization of the healthy gut PHI network.

Although PHILM currently infers interactions at the genus level—primarily due to sample size constraints of existing datasets—this limitation does not appear to hinder its utility for disease prediction. In our disease cohorts, microbial profiles were generated at the species level, and the lowest training sample-to-feature-ratio (*N*_train_/*N*_taxa_) was approximately 0.015, far below the conventional 1.0 threshold for reliable interaction inference. More importantly, PHILM-derived representations consistently outperform other relative abundance-based features in disease prediction, especially when both prokaryotic and phage profile-based Shannon diversities differ significantly between cases and controls. These results indicate that PHILM’s latent representations could capture subtle, predictive signals beyond simple taxonomic shifts and offer potential for non-invasive diagnosis across diverse diseases.

Despite its advantages, we acknowledge several limitations of the current framework. First, PHILM does not distinguish temperate from virulent phages—a distinction that may require genomic context or auxiliary sequencing data beyond abundance information. Second, although synthetic benchmarks yielded higher Pearson correlations than real-world datasets, real samples can be influenced by inter-prokaryotic interactions —which we showed in synthetic data can be incorporated into PHILM once a reliable interaction matrix becomes available—as well as nutrient gradients, host immune factors, and potential false-positive taxonomic assignments, all of which warrant further study. Third, the model is restricted to DNA phages and bulk metagenomic data, suggesting its extension to RNA phages or virus-like particle (VLP) metagenomes will depend on the availability of sufficient metatranscriptomic and VLP enrichment datasets. Finally, because our predictions operate at the genus level, larger and more targeted sample collections, especially from specific populations or disease states, will be required to achieve finer taxonomic resolution.

In summary, by leveraging the vast repositories of existing metagenomic data, PHILM provides a scalable, mechanism-agnostic approach to PHI inference. With appropriate reference databases, this framework can be extended to microbiota from other human body sites or environments. PHILM offers an efficient and robust alternative to traditional assembly- and co-abundance-based methods for reconstructing PHIs and predicting host metadata. Its flexible architecture also holds promise for broader applications in other cross-domain interaction predictions.

## Methods

### Synthetic data generated using ecological models

#### Generalized Lotka-Volterra model

We first generated synthetic data for a pool containing *N* prokaryotes and *M* phages using generalized Lotka-Volterra (GLV) population dynamics. We defined the GLV model as below^47,48^:

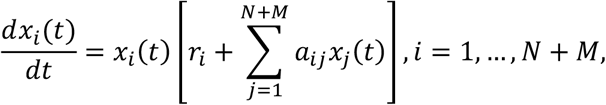

where *x*_*i*_(*t*) denotes the abundance of the *i*-th prokaryote/phage at time *t* ≥ 0. The GLV model has two major parameters: the interaction matrix *A* = (*a*_*ij*_) ∈ ℝ^(*N*+*M*)×(*N*+*M*)^, and the intrinsic growth-rate vector *r* = (*r*_*i*_) ∈ ℝ^(*N*+*M*)^. The parameter *a*_*ij*_ denotes the interaction strength of microorganism *i* to the per-capita growth rate of microorganism *j*. The parameter *r*_*i*_ is the intrinsic growth rate of microorganism *i*. The prokaryote-prokaryote interaction strengths were randomly choosing from 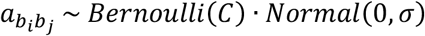 if *b*_*i*_ ≠ *b*_*j*_ and 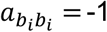, where *C* = 0.1 and σ = 0.05. We assume each prokaryotic host can be targeted by 10 phages and the interaction from prokaryotes to phages were selected from 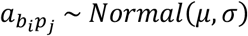 where *μ* = 0.9 and σ = 0.1. The interaction from phages to prokaryotes were selected from 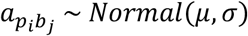 where σ = 0.1, *μ* = −0.5 for 20% of the phages, *μ* = 0.5 for the remaining 80%. The GLV model is designed to reflect the narrow host ranges of phages. We assume extremely strong inter-phage competition, such that when sampling from the phage pool, only a single and randomly selected phage taxon can infect a given prokaryotic host. From a pool consisting of *N* = 100 prokaryotes and *M* = 1,000 phages, we generated 200 to 1,800 samples in steps of 100 using the GLV model, with each pair of samples sharing 80% of prokaryotes. After confirming that all microorganisms had reached steady state at a simulation time of T = 1000, we extracted paired phage and prokaryotic profiles from each sample.

To evaluate the impact of inter-prokaryotic interactions, we generated between 200 and 1,800 synthetic samples by setting the GLV connectivity parameter to *C* = 0.9 and the interaction-strength standard deviation to σ = 0.25, while holding all other parameters constant.

#### Dynamic population model

Next, we generated synthetic data for a pool comprising N prokaryotes and M phages by employing a more sophisticated dynamic population (DP) model, adapted from the “more comprehensive model including sensitive prokaryotes (namely the S4 model)” proposed by Kimchi et al.^49^. Compared to the GLV model, our DP model resembles a Susceptible-Infected-Recovered (SIR) model in epidemiology and considers a variety of phage-host interaction processes including infection, lysis/lysogeny, induction, reversion and phage death. This model also permits a prokaryotic host that is already infected by a certain phage to acquire a second infection by another phage, yet it cannot be infected by a third one. In detail, the population densities of sensitive prokaryotes *S*_*i*_, phage *P*_*j*_ and their associated lysogens *L*_*ij*_ change according to the following three equations, respectively:

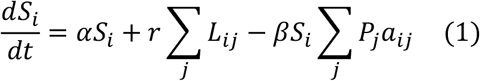

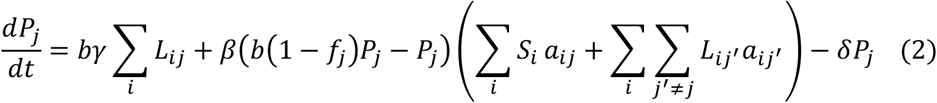

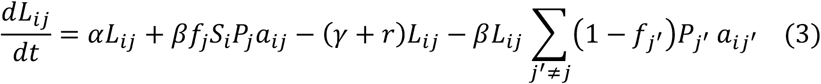

Here, *A* = (*a*_*ij*_) ∈ ℝ^(*N*+*M*)×(*N*+*M*)^ is a pre-defined binary matrix denoting if phage *j* is capable to infect prokaryote *i*, where we assume that a prokaryotic host can be infected by multiple phage predators and a phage can also target multiple prokaryotic hosts. α indicates the prokaryotic growth rate, while β indicates the phage infectivity. γ represents the prophage induction rate and δ represents the phage death rate. *b* denotes the burst size, the number of new virus particles released from a single infected host cell when it is lysed. *r* is defined as the reversion rate for a lysogen to lose a prophage. *f* is the fraction of phages’ infections leading to lysogeny, where phages with *f* = 0 are obligate lytic and have no associated lysogens. *f* is randomly chosen for each phage from a list of [0.0, 0.2, 0.4, 0.6, 0.8], representing different phage lysogeny abilities. Details and literature support for the parameter settings are provided in **Supplementary Table 7**. Equation (1) defines that the population density of *i*-th uninfected prokaryote is increased by self-growth and reversion from infected prokaryotes *L*_*ij*_, while decreased by the infection of phage *j* if their interaction is set to non-zero in the matrix *A*. Equation (2) defines that the population density of *j* -th phage is positively affected by the new phage particles produced by the burst from prophage induction and from the direct lysis of sensitive prokaryotes and those infected by other phages. The density is negatively affected by the phage infection and integration into available prokaryotic genomes (including sensitive prokaryotes and prokaryotes infected by other phages) and its death. Equation (3) defines the population density of the *i*-th prokaryote infected and integrated by the *j*-th phage in its genome (also known as lysogens). The density increases due to self-prokaryotic growth and phage *j* infecting sensitive prokaryote *i*, while it decreases due to phage induction, its conversion into a sensitive prokaryote and lysis by other phages *j*′ (where *j*′ ≠ *j*). From a pool comprising *N* = 100 prokaryotes and *M* = 1,000 phages, we generated between 200 and 1800 samples in increments of 100 using the DP model, with each sample containing exactly 10 prokaryotes, and each pair of samples shared 60% of prokaryotes. In each round of sampling, a random number of phages was selected for a specific prokaryote. After simulating for a longer period (T=5,000 steps) than the GLV model, instead of reaching a steady state, the phage and prokaryotic abundances may still fluctuate in a chaotic way but maintain a dynamic balance. In fact, this chaotic dynamic population model was first proposed to explain the coexistence of lytic and temperate phages in nature^49^.

### Metagenomic sample collection and processing

#### Healthy individual dataset

These datasets include 7,016 metagenomic stool samples of sequencing depth greater than 1 million reads from healthy individuals ranging in age from 8-75 years old without antibiotics usage. The sample accessions were collected from curatedMetagenomicData^55^ and downloaded from the Sequence Read Archive (SRA) database. According to previously reported criterion, age data were grouped into five categories including children adolescents (8 ≤ age ≤ 18), young adult (18 < age ≤ 35), middle aged (35 < age ≤ 50), senior (50 < age ≤ 65), and elderly (age > 65). Metadata can be found in **Supplementary Table 5**.

#### Disease datasets

Human fecal raw metagenomic sequencing data and clinical metadata of the samples for (1) IBD and controls were collected and downloaded from the Inflammatory Bowel Disease Multi’omics Database (IBDMDB) (https://ibdmdb.org/results), including HMP2 and HMP2 Pilot studies^69,70^, PRJEB15371^71^, PRJNA1086048^72^, PRJNA400072^73^ and PRJNA429990^74^. (2) CRC and controls were from PRJDB4176^75^, PRJEB10878^76^, PRJEB12449^77^, PRJEB27928^78^, PRJEB6070^79^, PRJEB7774^80^, PRJNA389927^81^, PRJNA429097^82^, PRJNA447983^83^, PRJNA531273 (with PRJNA397112 as controls)^84^, PRJNA731589^85^, PRJNA763023^86^ and PRJNA936589 (downloaded from https://www.ncbi.nlm.nih.gov/bioproject/PRJNA936589/ since no paper was published). The metadata of CRC were partially organized from Zun et al.^87^ (3) Multiple sclerosis (MS) and controls were from PRJEB51635 (patients) and PRJEB41786/PRJEB41787 (controls)^88^. (4) Parkinson’s disease (PD) and controls were from PRJEB53403 and PRJEB53401^89^ and PRJNA834801^90^. (5) Liver cirrhosis (LC) and controls were obtained from PRJEB6337^91^, PRJNA373901^92^ and from a paired set where patients are from PRJNA912122 and controls from PRJNA838648^93^. (6) Urinary tract infection (UTI) and controls were from PRJNA400628^94^. (7) Irritable bowel syndrome (IBS) and controls were collected from PRJEB37924^95^. (8) COVID19 and controls were obtained from PRJDB13214^96^ and PRJNA650244^97^. (9) HIV and controls were obtained from PRJNA307231^98^, PRJNA692830^99^, PRJNA391226^100^, and PRJNA450025^101^. (10) Hypertension (HTN) and controls were from PRJEB13870^102^. More details in the metadata can be available in **Supplementary Table 8**.

Quality control and host contamination removal were done with KneadData v.0.12.0 (https://github.com/biobakery/kneaddata) using Trimmomatic v.0.39^103^ with options “ILLUMINACLIP:<adapter file>:2:30:10:2:TRUE LEADING:3 TRAILING:3 SLIDINGWINDOW:4:15 MINLEN:30” and Bowtie2 v.2.4.5^104^, respectively. Phanta v.1.1.0 was subsequently using the following parameters: “confidence_threshold: 0.1; class_mem_mb: 60000; class_threads: 28” and the prophage-masked version of database^32^. Specifically, to further filter false-positive species, different viral coverage thresholds and prokaryotic coverage thresholds have been selected based on the sample sequencing depths following the suggestions at https://github.com/bhattlab/phanta/tree/main. Phage and prokaryotic profiles were derived from the Phanta-generated relative taxonomic profiles. To only include viruses infecting prokaryotic hosts (namely phages), only viral taxa belonging to the orders *Caudovirales, Tubulavirales, Petitvirales, Cremevirales*, and *Halopanivirales* were considered.

### Deep learning model architectures and hyperparameter tuning

We evaluated and compared three deep learning models: Transformer^105^, Residual Neural Network (ResNet)^106^ and Neural Ordinary Differential Equations (NODE)^107^. We included the transformer because of its robust attention mechanism and its ability to capture long-range dependencies within the data^105^. ResNet was selected due to its use of skip connections, which facilitate the training of deeper architectures by ensuring direct gradient flow and mitigating the vanishing gradient problem^106^. NODE extends its precursor, ResNet, by integrating skip connections that are analogous to Euler steps in ODE solvers^107^. Each residual block in NODE represents a small time change of variables, inspiring its continuous-depth architecture. Unlike ResNet’s discrete hidden layers, NODE parameterizes the derivative of the hidden state using a neural network and computes outputs through an ODE solver. This methodology allows NODE to maintain constant memory, adapt its evaluation strategy for each input, and balance numerical precision with speed.

#### phiNODE

phiNODE is a method that integrates an implicit NODE module within a deep learning architecture. Prior to modeling, the centered log-ratio transformation is applied to both the phage-excluded and phage taxonomic abundance profiles. Considering the large disparity in the numbers of prokaryotic and phage taxa, it is not possible to directly apply the original NODE architecture^107^ as it requires input and output dimensions to be equal. Inspired by the architecture of mNODE^44^, we incorporated two fully connected hidden layers with the NODE in between to transform the data dimensions. The NODE module in the middle of the architecture computes the time evolution of ODEs whose first-order time derivatives are approximated by two fully connected layers. We used the NODE to reduce memory and training time as NODE shares connection weights across continuous layers and utilizes fewer parameters.

The architecture comprises three sequential modules: (1) A fully connected layer maps the input phage compositions to a hidden representation of dimension *N*_*h*_, followed by the hyperbolic tangent (tanh) activation function, (2) a neural ODE module approximates the first-order time derivative using a two-layer multilayer perceptron (MLP) (each layer with dimension *N*_*h*_) to implement the NODE^107^, with each layer followed by the tanh activation function, and (3) a fully connected layer transforms the hidden representation to yield the phage-excluded output profiles. The Adam optimizer^108^ is used for the gradient descent. To prevent overfitting, we used (1) the L2 regularization with the weight decay parameter λ in the Adam optimizer, (2) early stopping in the training. Training stops if the validation mean squared error (MSE) loss does not decrease by at least 1×10^−4^ or by at least 1% relative to the best observed loss over 20 consecutive epochs. Three hyperparameters are tuned based on the validation loss: the dimension of the hidden layer *N*_*h*_, the L2 regularization parameter λ and the learning rate *l*. Specifically, *N*_*h*_ is selected from the three candidate values closest to the number of output features within the set [4, 8, 16, 32, 64, 128, 256, 320, 512, 640, 1024, 2048, 2560, 4096], λ is chosen from the range of 10^−6^ to 10^−2^ and *l* is selected from 10^−8^ to 10^−5^. The optimal hyperparameter combination was determined using the Optuna v.4.2.1 framework in 10 trials^109^.

To enable the model to explicitly consider inter-prokaryotic interactions, we appended a graph convolutional network (GCN) layer defined by

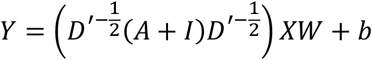

where *X* ∈ ℝ^*B*×*N*^ is the batch of phiNODE originally predicted prokaryotic abundances (*B*: batch size, *N*: taxa), *A* ∈ ℝ^*N*×*N*^ is the normalized Laplacian of the inter-prokaryotic interaction network, and *I* is the identity matrix that adds self-loops. The diagonal matrix *D*^i^ has entries 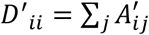, so that 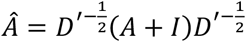 symmetrically normalizes the graph. The smoothed feature matrix *AX* is mapped back to the output space by the learnable weight *W* ∈ ℝ^*N*×*N*^ and bias *b* ∈ ℝ^*N*^, yielding *Y* ∈ ℝ^*B*×*N*^, which is the final inter-prokaryotic interaction-aware profile predictions. This operation ensures that each taxon’s final prediction is informed by both its own abundance and those of its interacting partners.

#### phiResNet

phiResNet adopts the residual neural network architecture that learns complex representations while mitigating the vanishing gradient problem. The architecture is organized into three sequential modules: (1) An input layer maps the input phage compositions to a hidden representation of dimension *N*_*h*_, (2) a series of hidden layers, each implemented as a fully connected layer, is arranged with residual connections: the output of each hidden layer is added to its input, and the sum is passed through a ReLU activation function, and (3) an output layer projects the final hidden representation to the output dimensionality. Hyperparameter tuning (the dimension of the hidden layer *N*_*h*_, the L2 regularization parameter λ and the learning rate *l* and early stopping was implemented using the same strategy as that employed for phiNODE.

#### phiTransformer

phiTransformer, based on the transformer architecture, is designed to capture contextual dependencies in input data through self-attention mechanisms. Its architecture is composed of the following modules: (1) a linear layer projects the input features from their original dimension to a higher-dimensional space *d*_*model*_, where the embedded input is then reshaped (by adding an extra dimension) for compatibility with subsequent layers, (2) a multi-layer transformer encoder (layer numbers determined by *N*_layer_), constructed from identical encoder layers, that applies multi-head self-attention and feedforward operations (with dropout regularization *r*_*dropout*_) to extract rich representations from the embedded inputs, (3) a corresponding multi-layer transformer decoder (layer numbers determined by *N*_*layer*_) that takes a target tensor (initialized to zeros) along with the encoder’s output (keys and values) to generate context-aware features with each decoder layer uses multi-head attention (*N*_*head*_ = max(2, int(*d*_*model*_/8))) and feedforward networks (*d*_*feedforward*_ = 2 · *d*_*model*_), and (4) a linear layer transforms the decoder’s output into the final output with the desired dimensionality. To reinforce the original signal and stabilize training, a residual connection is implemented by linearly mapping the original input directly to the output and summing it with the decoder’s result. A total of five hyperparameter are tuned based on the validation loss: for the model dimension *d*_*model*_, the number of encoder/decoder layers *N*_*layer*_, the dropout rate *r*_*dropout*_, the L2 regularization parameter λ and the learning rate *l*. Specifically, *d*_*model*_ is selected from the three candidate values closest to the number of output features within the set [4, 8, 16, 32, 64, 128, 256, 320, 512, 640, 1024, 2048, 2560, 4096], *N*_*layer*_ is chosen from [2, 4, 6, 8], *r*_*dropout*_ is selected from [0.3, 0.4, 0.5], while other hyperparameters were tuned following a strategy analogous to that used for phiNODE. Model construction, training and validation are implemented using PyTorch v2.4.1^110^.

### Model training, validation and testing

For synthetic and real-world datasets, deep learning models including phiNODE, phiResNet and phiTransformer were trained on 80% of samples (as the training set). Their hyperparameters were heuristically determined on 10% of the samples (as the validation set), and the mean PCCs of all prokaryotic taxa across samples on the remaining 10% samples (as the test set) were calculated to evaluate the models’ abilities to predict prokaryotic compositions.

### Inferring phage-host interactions via sensitivity analysis

The well-trained deep learning model within PHILM takes the relative abundance of phage *i* (*x*_*i*_) as inputs and generates predictions for the relative abundance of prokaryote *j* (y_*j*_). For the sample *m* in the training set, 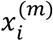 for all *i* is provided as the input vector to the trained model and the model can predict the relative abundances 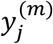 for all prokaryotes. To investigate the influence of phage *i* on prokaryote *j* for a particular sample *m*, we perturb 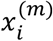 by setting it as the mean of relative abundances for phage *i* across samples (denoted as 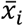) while keeping values of 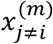 intact. Thus, the perturbation amount for phage *i* is 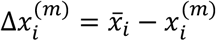. This perturbed vector for phage relative abundances is provided to the trained deep learning model to regenerate predictions for prokaryotic relative abundance 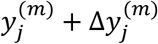 where 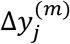 is the deviation of newly predicted relative abundance for prokaryote *j* when the deep learning model uses the perturbed input vector from that when it uses the unperturbed input vector. For the sample *m*, the sensitivity of prokaryote *j* to phage *i* can be defined as 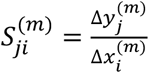. Sensitivity value is averaged across samples and z-score normalized for each phage. The overall sensitivity of prokaryote *j* to phage *i* is 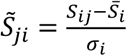, where 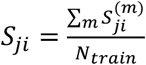 and *N*_*train*_ is the number of samples in the training set. 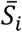and σ_*i*_ are the mean and standard deviation, respectively, of *S*_*ji*_ calculated for phage *j*.

### Network analysis and visualization

The phage-host interaction network was visualized using Cytoscape v.3.10.3^111^. Modules were identified using the community clustering algorithm GLay^112^, which is available as an app within Cytoscape.

### Phage-host interactions at the genus level in the Metagenomic Gut Virus catalogue

The phage-host associations in the Metagenomic Gut Virus (MGV) catalogue were sourced from https://portal.nersc.gov/MGV/MGV_v1.0_2021_07_08/mgv_host_assignments.tsv (accessed on January 24th, 2025). In this dataset, host genomes from the Unified Human Gastrointestinal Genome (UHGG) database^113^ were linked to viruses via a combination of CRISPR spacer matches and near-identical 1 kb BLAST hits between phage and prokaryotic genomes^36^. As the taxonomy for prokaryotic hosts is based on the Genome Taxonomy Database (GTDB r89), we converted GTDB taxonomy to NCBI taxonomy at the genus level using gtdb_to_taxdump (https://github.com/nick-youngblut/gtdb_to_taxdump)^114^.

### Predict phage-host interactions using a co-abundance-based method

FastSpar v.1.0.0 was used to calculate the co-abundance correlations between the prokaryotic and phage taxa^40^. P-values were generated by performing 1000 random permutations. Statistically significant (P < 0.05) correlations with magnitude exceeding certain thresholds were considered as potential PHIs.

### Infer phage-host interactions using the assembly-based approach

For the 7,016 metagenomic stool samples from healthy individuals, we performed single-sample assembly using MEGAHIT v.1.2.9^115^, after excluding contigs less than 2,000 bp, resulting in a total of 64,888,908 assembled contigs. Viral and non-viral sequences were differentiated using geNomad v.1.7.6^27^, which excluded provirus and plasmid sequences, yielding 62,053,011 non-viral contigs and 2,835,897 viral contigs. Non-viral contigs were binned using MetaBAT v.2.15^116^, producing 206,411 bins, which were further assessed for quality using CheckM v.1.2.3^117^, retaining those with ≥50% completeness and <10% contamination, leading to 90,447 metagenome-assembled genomes (MAGs). Taxonomic classification of prokaryotic bins was performed using CAT pack v.6.0.1^118^ with HumGut as the reference database^119^, identifying 45 prokaryotic genera. CRISPR spacers were detected in the MAGs using CasCRISPRTyper v.1.8.0^120^, yielding 56,461 CRISPR spacers. Viral contigs were quality-filtered using CheckV v1.0.3^121^, retaining 96,599 high-to medium-quality viral contigs, which were then assigned taxa using uhgv-tools (https://github.com/snayfach/UHGV/blob/main/CLASSIFY.md) with MGV as the reference database^30^, identifying 1,947 viral genera. To predict host-phage interactions, we followed the strategy described by Nayfach et al.^56^ and employed CRISPR spacer matching via blastn-short (allowing ≤1 mismatch) and sequence similarity-based matching via BLASTN v.2.15.0+^122^, considering hits with >90% identity and >500 bp overlap. These approaches identified 2,589 host MAG-viral contig interactions, resolving into 221 host-virus interactions at the genus level.

### Machine learning classifiers

The machine learning models were trained and validated using a nested cross-validation framework to ensure robust model selection and performance evaluation. Input features were standardized using StandardScaler in the scikit-learn module^123^. Three classification models were evaluated: Support Vector Machine (SVM), Logistic Regression (LR) and Random Forest (RF). Model selection and performance assessment were conducted using a 5-fold cross-validation, with folds generated via StratifiedKFold from the scikit-learn library^123^. One of the five folds was reserved as a held-out test set. This approach was employed to address class imbalance in the data, ensuring that each fold preserved the original dataset’s class distribution and thereby providing a more robust evaluation of model performance. The remaining data were subjected to hyperparameter optimization using GridSearchCV from the scikit-learn library with an inner 5-fold cross-validation^123^. For SVM model, the hyperparameter grid included regularization values (C) of 0.1, 1, and 10 and kernel types of linear and radial basis function (rbf). For LR model, the hyperparameter grid explored different penalty types (“none”, L2, and ElasticNet) and regularization strength (C) values of 0.1, 1, and 10. The hyperparameter grid for the Random Forest included the number of trees (n_estimators: 100, 300, 500) and the fraction of samples used for building each tree (max_samples: 0.2, 0.5, 0.8). Model selection was based on the highest mean area under the curve (AUC) score obtained from the inner cross-validation, with the best hyperparameters applied to the outer test fold for evaluation. Model performance was assessed using AUC, precision, recall, and F1-score, and predictions for each fold. For age data, which was treated as a multi-class classification problem, we computed one-vs-all AUC as well as macro-averaged precision, recall, and F1-score.

## Acknowledgements

Y.-Y.L. acknowledges funding support from the Resnek Family Center for PSC Research at Brigham and Women’s Hospital, the National Institutes of Health (R01AI141529, R01HD093761, RF1AG067744, UH3OD023268, U19AI095219 and U01HL089856) as well as the Office of the Assistant Secretary of Defense for Health Affairs, through the Traumatic Brain Injury and Psychological Health Research Program (Focused Program Award) under award no. W81XWH-22-S-TBIPH2, endorsed by the Department of Defense. X.-W.W. acknowledges funding support from the National Institutes of Health (K25HL166208).

## Data and code availability

All code for running PHILM is available at https://github.com/YiyanYang0728/phiNODE. The scripts for data processing and figure generation can be found at https://doi.org/10.5281/zenodo.15500282.

## Notes

https://doi.org/10.5281/zenodo.15500282

### Competing Interest Statement

The authors have declared no competing interest.

### Summary of Updates

The manuscript has been revised to emphasize the overarching deep-learning framework rather than any particular model architecture within it. Main text, figures and SI have been modified accordingly too.

