## Supplementary Figures for "Deep Learning Transforms Phage-Host Interaction Discovery from Metagenomic Data"

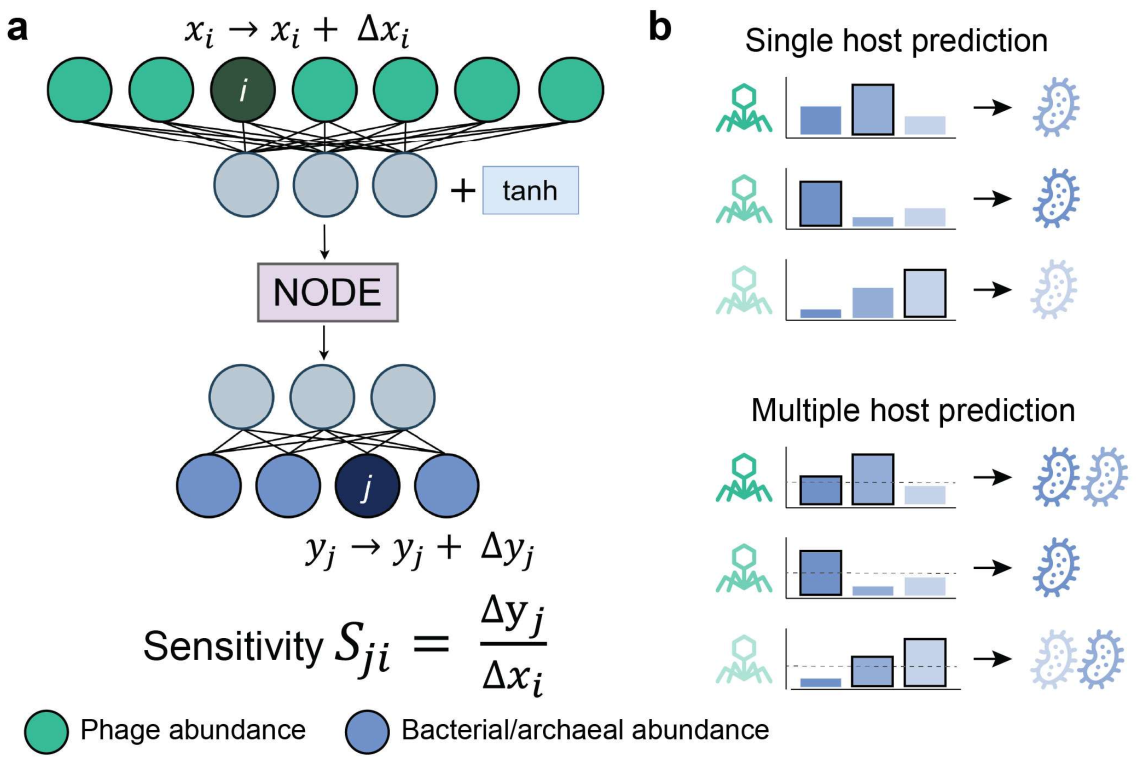


**Supplementary Figure 1**. Sensitivity analysis to predict putative hosts for phages. **a**, The sensitivity value of a prokaryotic taxon *j* to phage taxon *i*, denoted as $S_{ji}$, is defined as the ratio between the deviation in the relative abundance of prokaryotic taxa *j* ($\Delta y_j$) and the perturbation amount in the relative abundance of phage taxa *i* ($\Delta x_i$). The phiNODE architecture is shown here as an exemplar model. **b**, There are two host prediction modes to determine phage hosts. The single-host mode predicts the prokaryote *j* with the largest $S_{ji}$ as the putative host for phage *i*, while the multi-host mode predicts any prokaryote *j* with $S_{ji}$ greater than a threshold as the putative host.

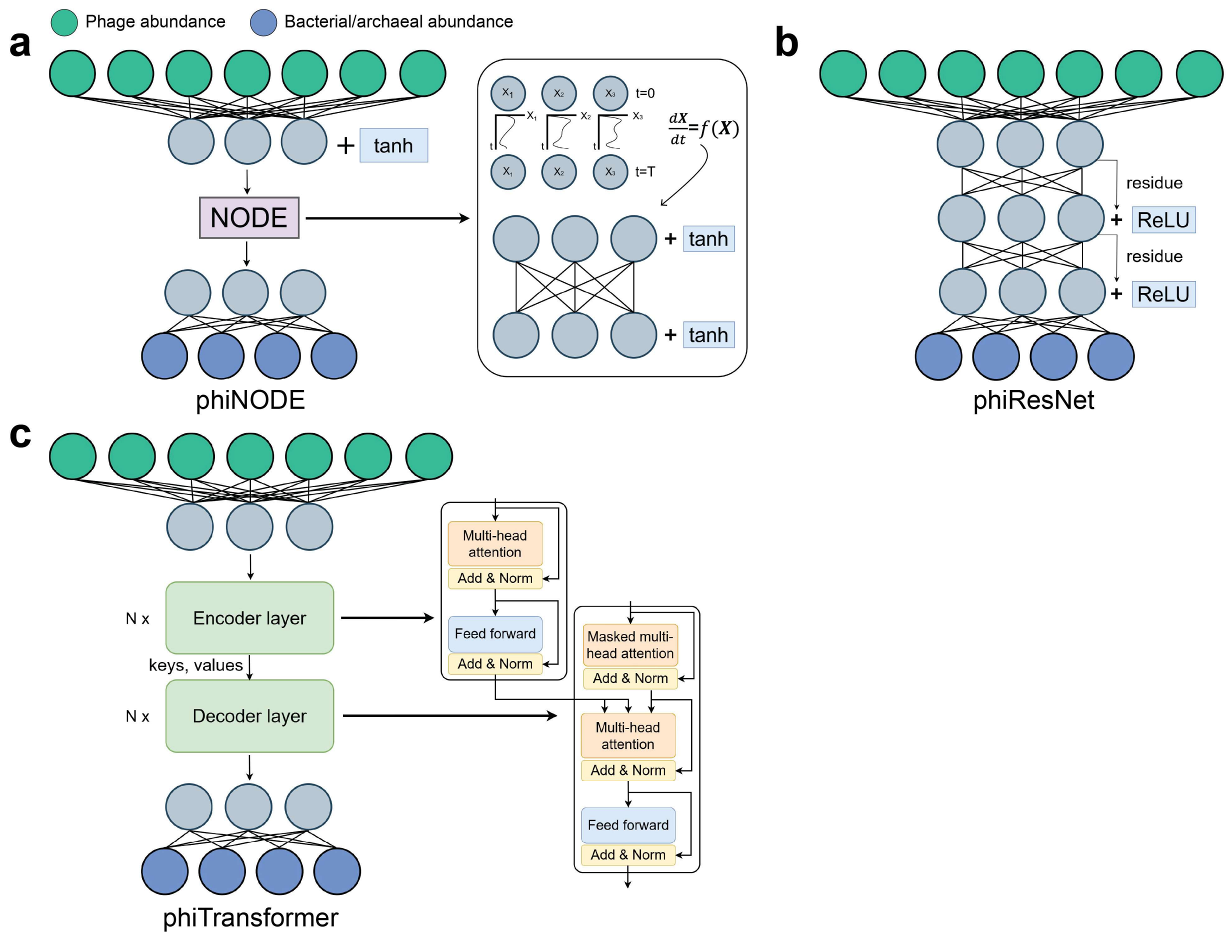


**Supplementary Figure 2**. Deep learning model architectures tested in PHILM. Grey nodes represent neurons in hidden layers in the models. **a**, The architecture of phiNODE. The NODE is a module in the middle of the architecture computes the time evolution of ODEs whose first-order time derivatives are approximated by two fully connected neural network layers. **b**, The architecture of phiResNet. It consists of three sequential modules: an input layer; a stack of fully connected hidden layers with residual connections—each hidden layer's output is added to its input and passed through a ReLU activation; and a final output layer. **c**, The architecture of phiTransformer. It comprises four modules: a linear layer, where the input is embedded and reshaped for compatibility with subsequent layers; a multi-layer transformer encoder that applies multi-head self-attention and feedforward operations to extract rich representations from the embedded inputs; a corresponding multi-layer transformer decoder that takes a target tensor along with the encoder's output (keys and values) to generate context-aware features using multi-head attention and feedforward networks; and a linear layer transforms the decoder's output into the final output.

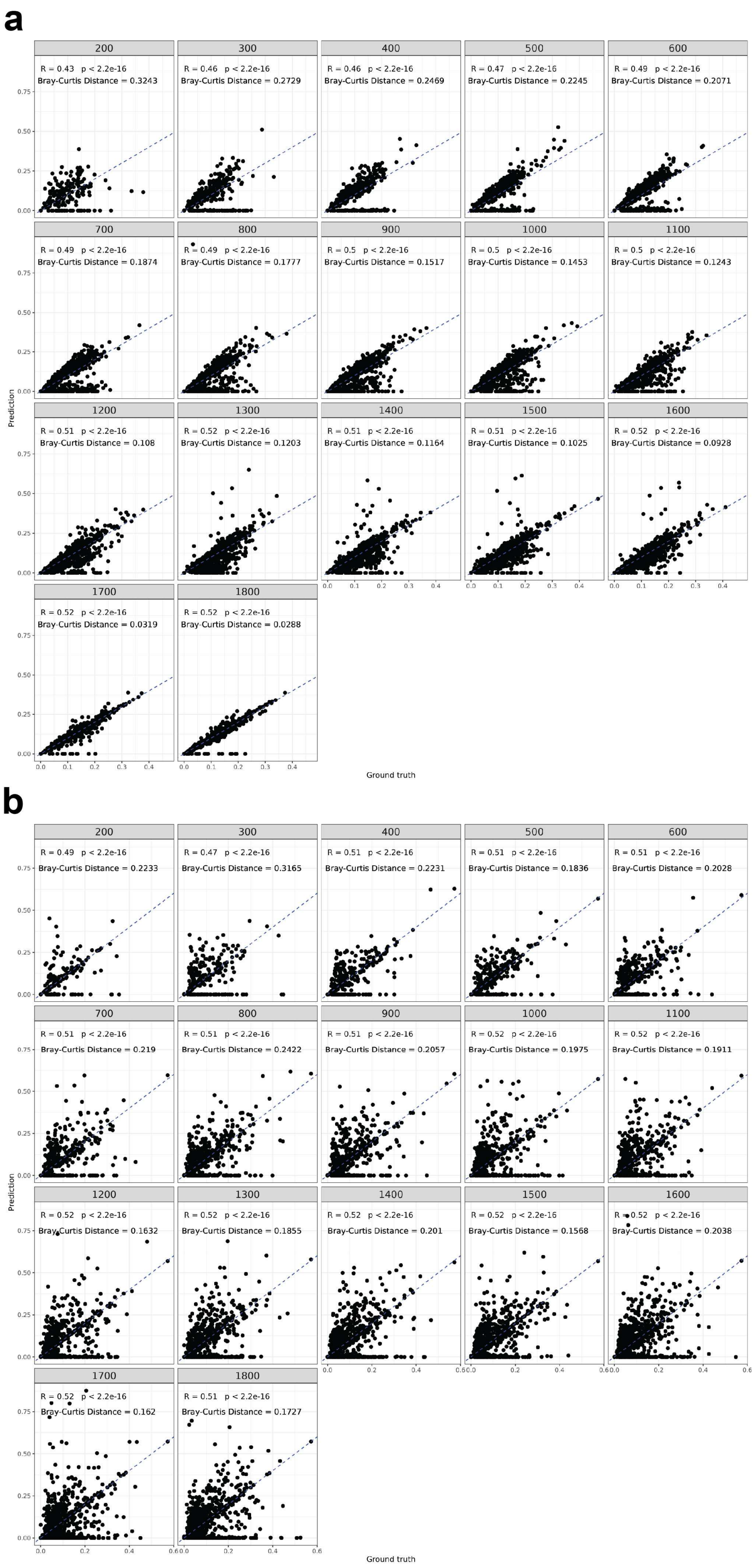
a
200
R = 0.43 p < 2.2e-16
Bray-Curtis Distance = 0.3243
300
R = 0.46 p < 2.2e-16
Bray-Curtis Distance = 0.2729
400
R = 0.46 p < 2.2e-16
Bray-Curtis Distance = 0.2469
500
R = 0.47 p < 2.2e-16
Bray-Curtis Distance = 0.2245
600
R = 0.49 p < 2.2e-16
Bray-Curtis Distance = 0.2071
700
R = 0.49 p < 2.2e-16
Bray-Curtis Distance = 0.1874
800
R = 0.49 p < 2.2e-16
Bray-Curtis Distance = 0.1777
900
R = 0.5 p < 2.2e-16
Bray-Curtis Distance = 0.1517
1000
R = 0.5 p < 2.2e-16
Bray-Curtis Distance = 0.1453
1100
R = 0.5 p < 2.2e-16
Bray-Curtis Distance = 0.1243
1200
R = 0.51 p < 2.2e-16
Bray-Curtis Distance = 0.108
1300
R = 0.52 p < 2.2e-16
Bray-Curtis Distance = 0.1203
1400
R = 0.51 p < 2.2e-16
Bray-Curtis Distance = 0.1164
1500
R = 0.51 p < 2.2e-16
Bray-Curtis Distance = 0.1025
1600
R = 0.52 p < 2.2e-16
Bray-Curtis Distance = 0.0928
1700
R = 0.52 p < 2.2e-16
Bray-Curtis Distance = 0.0319
1800
R = 0.52 p < 2.2e-16
Bray-Curtis Distance = 0.0288
Prediction
Ground truth
b
200
R = 0.49 p < 2.2e-16
Bray-Curtis Distance = 0.2233
300
R = 0.47 p < 2.2e-16
Bray-Curtis Distance = 0.3165
400
R = 0.51 p < 2.2e-16
Bray-Curtis Distance = 0.2231
500
R = 0.51 p < 2.2e-16
Bray-Curtis Distance = 0.1836
600
R = 0.51 p < 2.2e-16
Bray-Curtis Distance = 0.2028
700
R = 0.51 p < 2.2e-16
Bray-Curtis Distance = 0.219
800
R = 0.51 p < 2.2e-16
Bray-Curtis Distance = 0.2422
900
R = 0.51 p < 2.2e-16
Bray-Curtis Distance = 0.2057
1000
R = 0.52 p < 2.2e-16
Bray-Curtis Distance = 0.1975
1100
R = 0.52 p < 2.2e-16
Bray-Curtis Distance = 0.1911
1200
R = 0.52 p < 2.2e-16
Bray-Curtis Distance = 0.1632
1300
R = 0.52 p < 2.2e-16
Bray-Curtis Distance = 0.1855
1400
R = 0.52 p < 2.2e-16
Bray-Curtis Distance = 0.201
1500
R = 0.52 p < 2.2e-16
Bray-Curtis Distance = 0.1568
1600
R = 0.52 p < 2.2e-16
Bray-Curtis Distance = 0.2038
1700
R = 0.52 p < 2.2e-16
Bray-Curtis Distance = 0.162
1800
R = 0.51 p < 2.2e-16
Bray-Curtis Distance = 0.1727
Prediction
Ground truth

**Supplementary Figure 3**. Comparison of predicted versus ground-truth prokaryotic relative abundance profiles on synthetic testing datasets generated by (**a**) the GLV model and (**b**) the DP model. X-axis represents predicted prokaryotic relative abundance in a sample, while y-axis represents the true abundance. Each small panel corresponds to a different sample size, plotting predicted relative abundances against ground truth values. The blue dashed diagonal line indicates perfect agreement. For each sample size, the Spearman correlation coefficient (R), its associated p-value, and the Bray-Curtis distance between predicted and true values are reported.

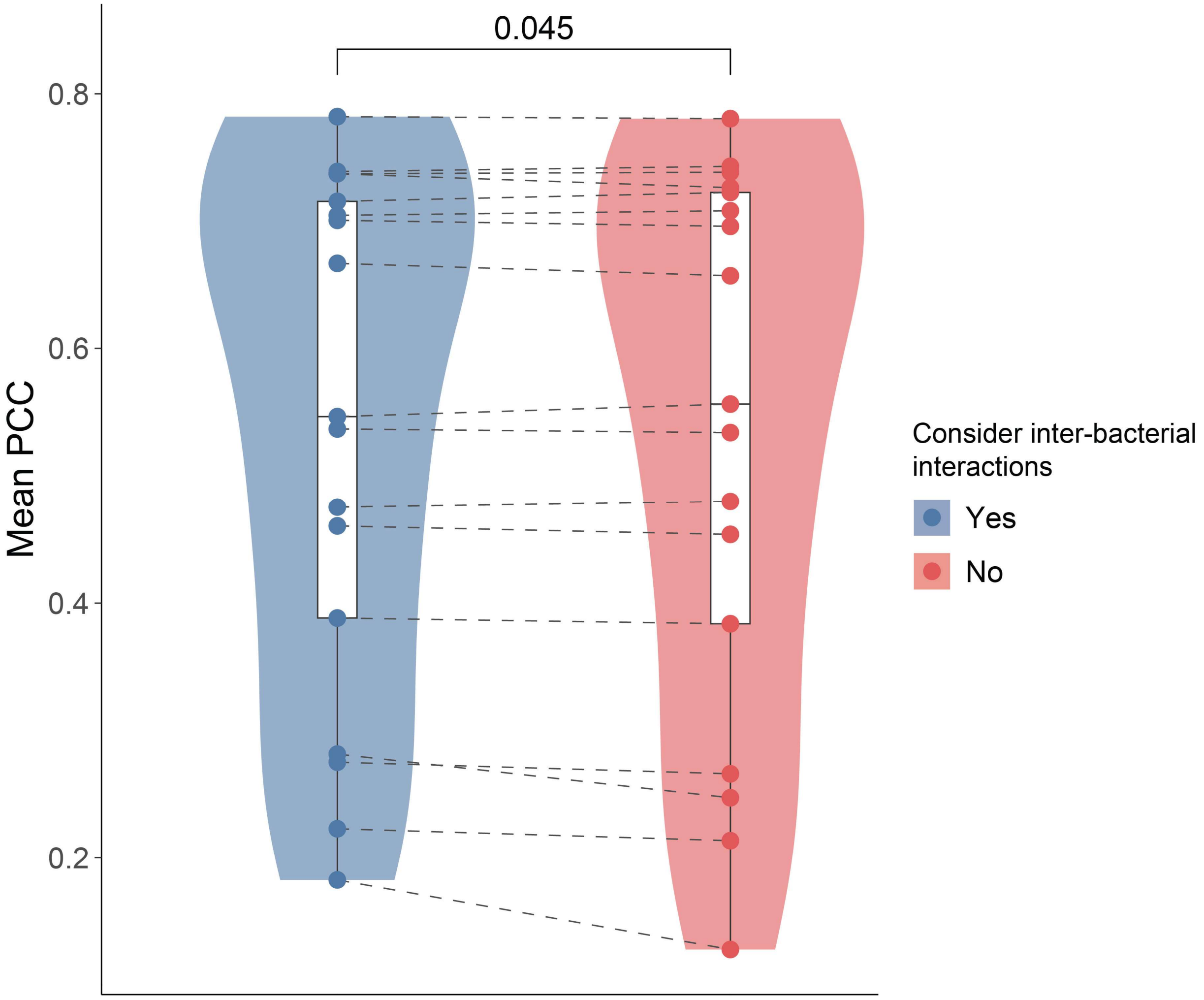


**Supplementary Figure 4**. Comparison of mean PCC values obtained by PHILM (based on phiNODE model) on GLV-generated test data using loss functions that either account for inter-bacterial interactions (blue) or ignore them (red). Paired dots (connected by lines) show the paired mean PCC for each training sample size. The p-value was calculated by a one-sided paired Student's t-test ($H_1$: inter-bacteria interaction-aware > inter-bacterial interaction-naive).

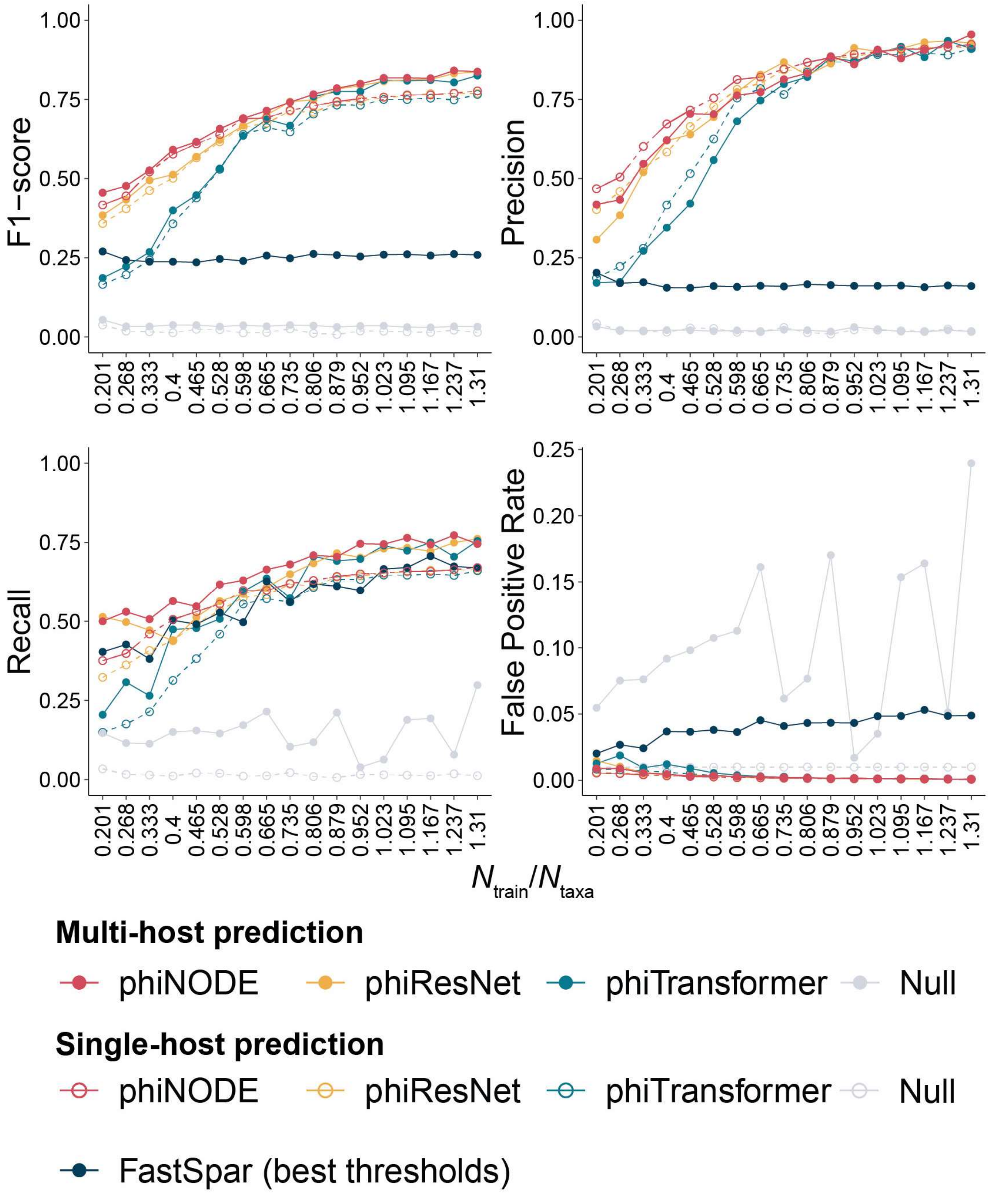


**Supplementary Figure 5.** F1-scores, precisions, recalls, and false positive rates for phiNODE, phiResNet, phiTransformer and FastSpar (using the optimal threshold that maximizes F1 scores), on synthetic data generated by DP model across varying $N_{train}/N_{taxa}$ ratios. Both single-host (dashed lines and empty circles) and multi-host (solid lines and filled circles) prediction modes were performed.

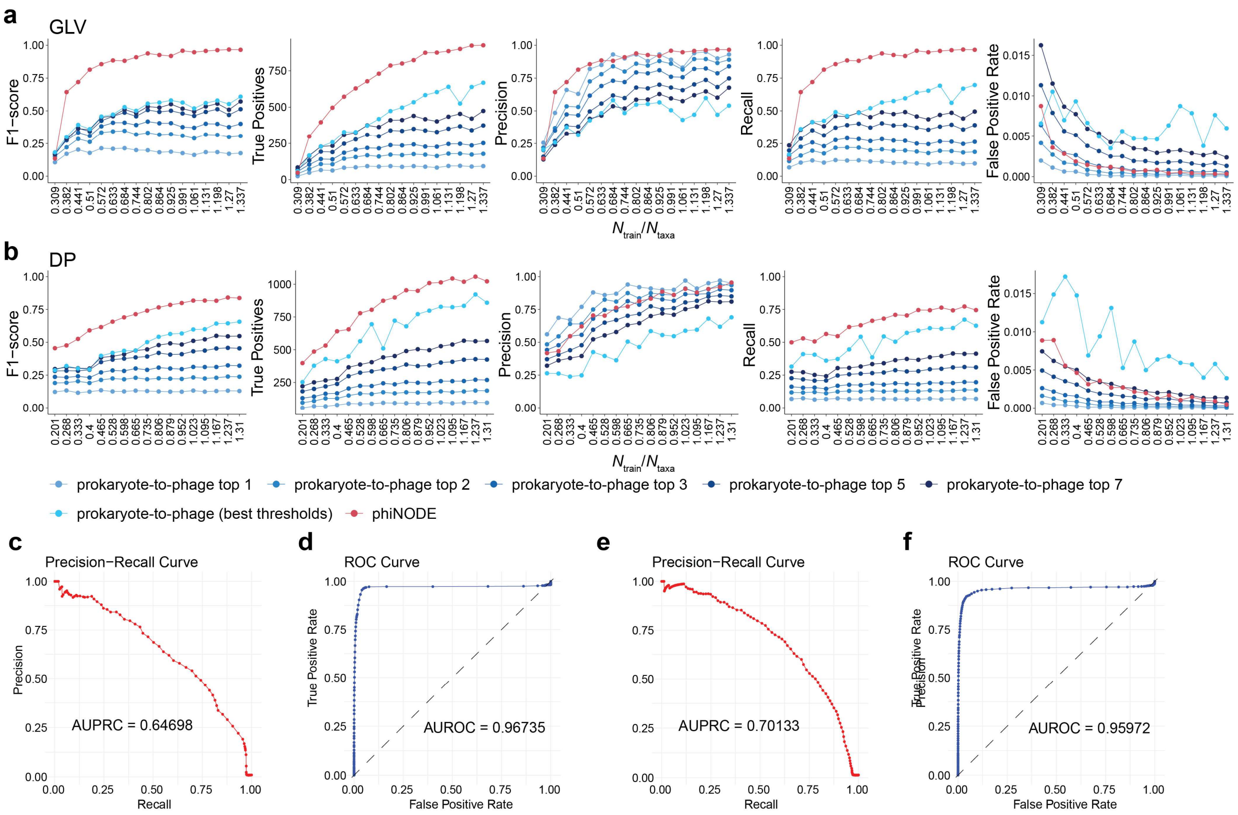


**Supplementary Figure 6**. Performance of prokaryote-to-phage method in predicting phage profiles from prokaryotic profiles (using the NODE model architecture) with different numbers of top phages and the optimal number of phages that maximizes F1-scores for inferring phage-host interactions in synthetic data from (**a**) GLV and (**b**) DP models across varying $N_{\text{train}}/N_{\text{taxa}}$ ratios. **c-f**, The (**c**,**e**) precision-recall and (**d**,**f**) receiver operating characteristic (ROC) curves for "Bac2phage" method applied to synthetic data with the largest sample size (N=1800) generated by the (**c-d**) GLV model and (**e-f**) DP model. Of note, here phiNODE used single host prediction for GLV model and multiple host prediction for DP model.

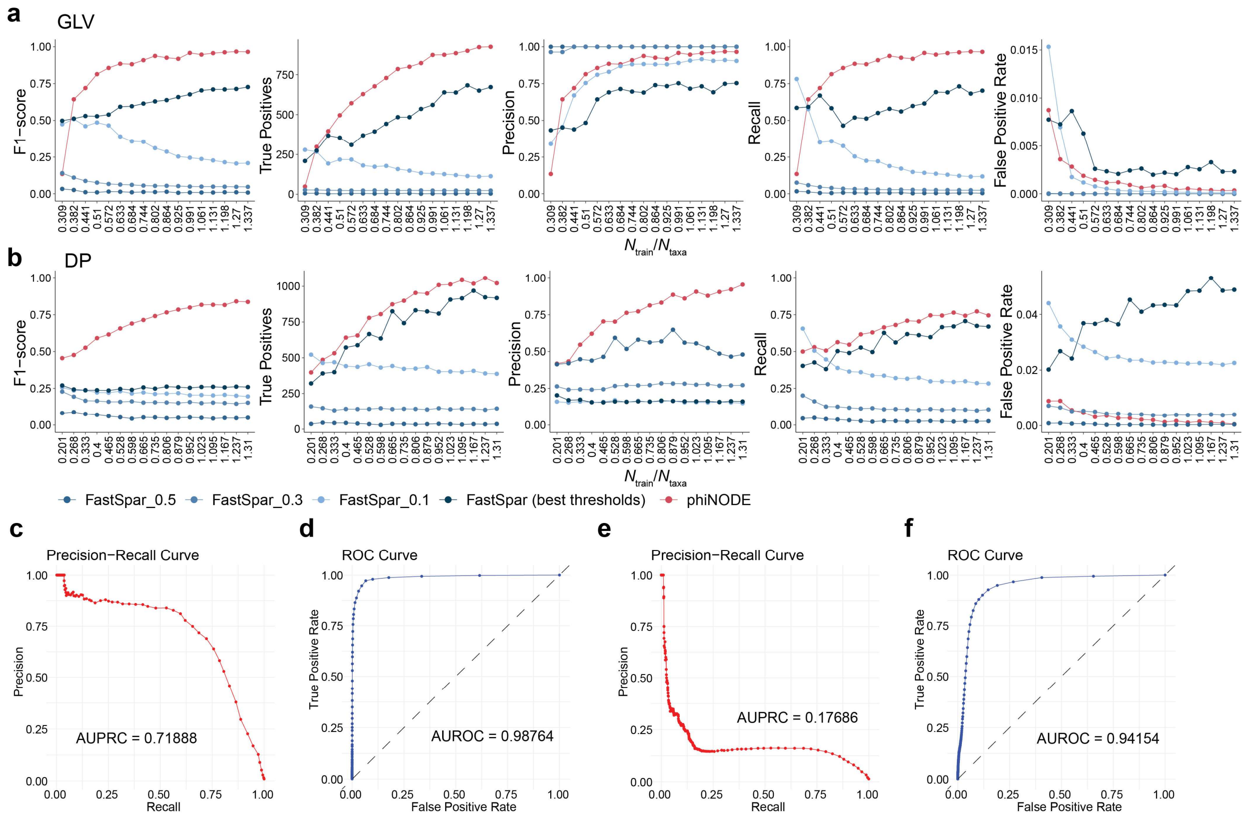


**Supplementary Figure 7**. Performance of FastSpar using thresholds of ±0.5, ±0.3, and ±0.1, and the optimal threshold maximizing F1-scores, for predicting phage-host interactions in synthetic data from (**a**) GLV and (**b**) DP models across varying $N_{train}/N_{taxa}$ ratios. For comparison, phiNODE single-host and multi-host predictions are also shown. **c-f**, The (**c**,**e**) precision-recall and (**d**,**f**) receiver operating characteristic (ROC) curves for FastSpar applied to largest sample size (N=1800) synthetic data with the largest sample size generated by the (**c-d**) GLV model and (**e-f**) DP model. Of note, here phiNODE used single host prediction for GLV model and multiple host prediction for DP model.

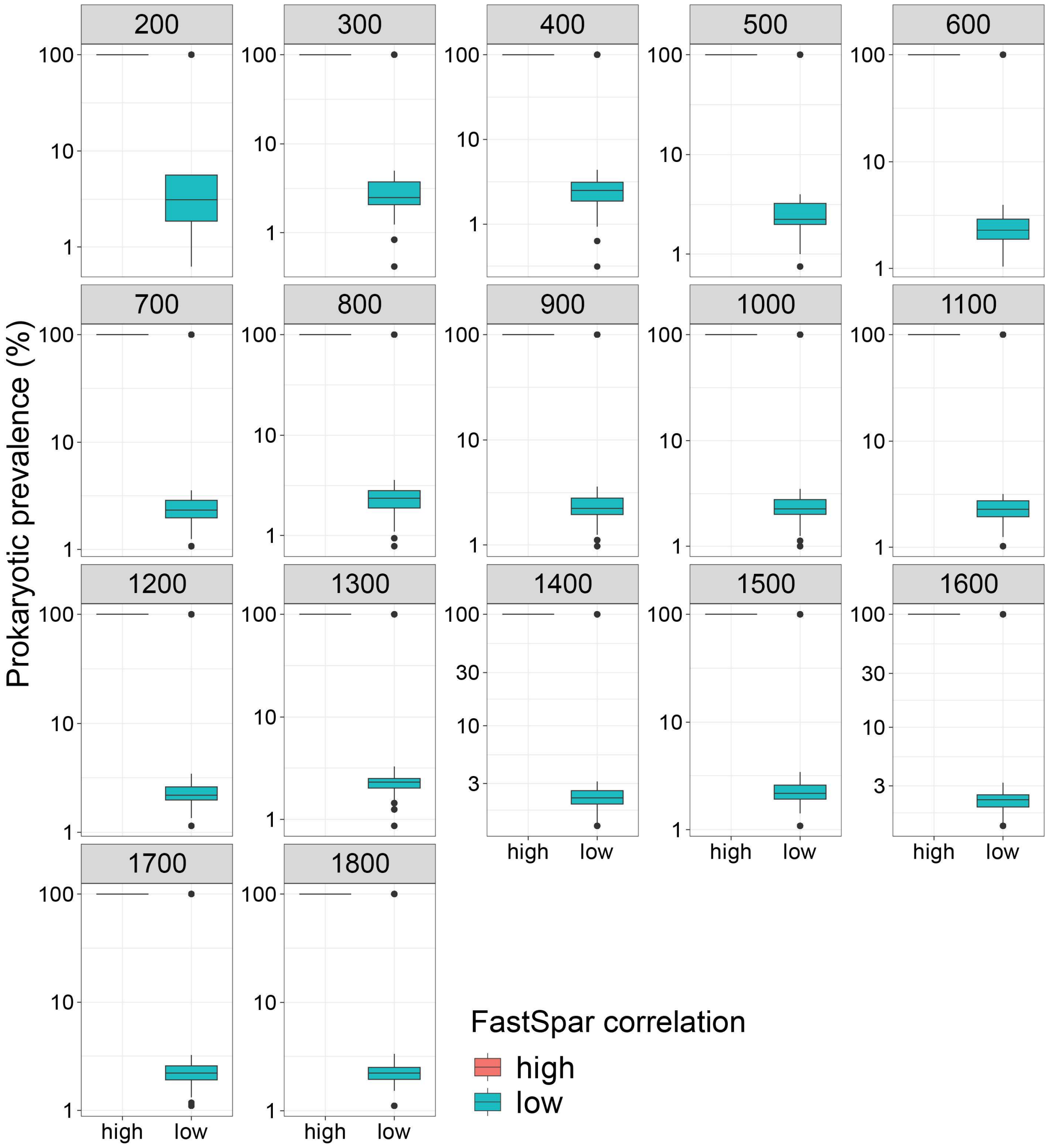


**Supplementary Figure 8**. FastSpar tends to assign high correlations to interactions involving prevalent prokaryotes. Boxplots show the prokaryotic prevalence for phage-host interactions with high and low FastSpar correlation values in synthetic data generated by the GLV model across different sample sizes. Interactions with FastSpar correlations greater than 0.5 or less than -0.5 are classified as high, while those with values in between are classified as low.

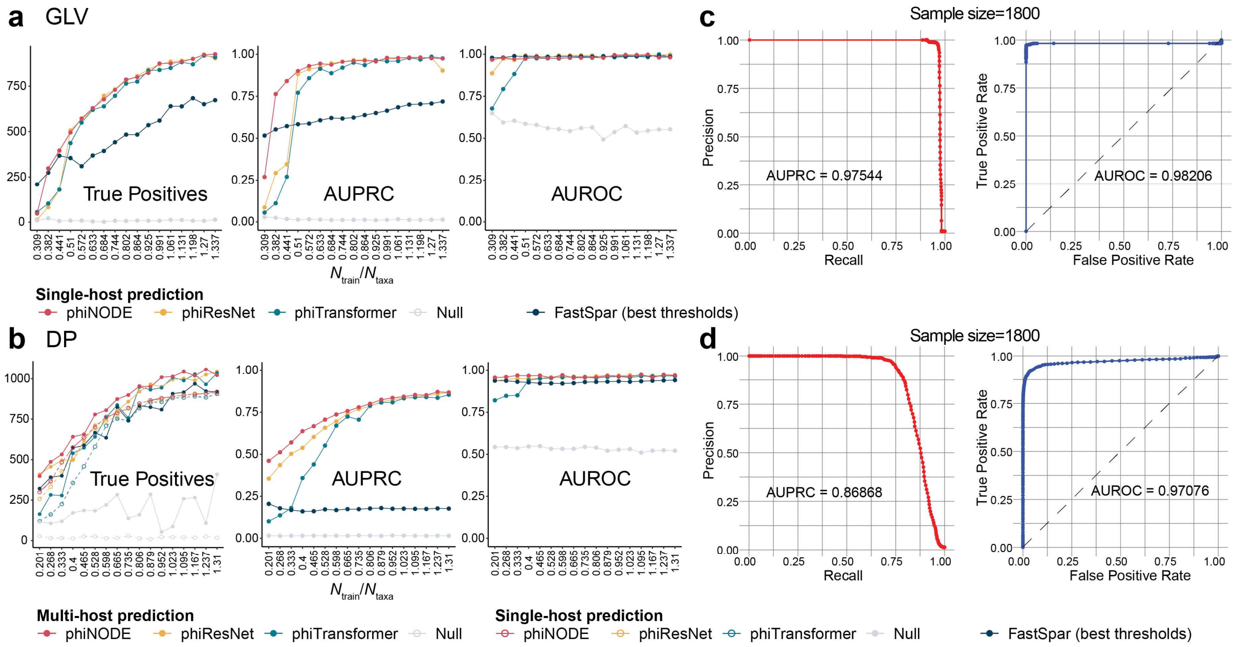


**Supplementary Figure 9**. True positives, the areas under the precision-recall curve (AUPRCs) and the areas under the receiver operating characteristic curve (AUROCs) for phiNODE, phiResNet, phiTransformer, and FastSpar (using optimal thresholds that maximize F1 scores) on synthetic data generated by (**a**) the GLV model and (**b**) the DP model, evaluated across varying $N_{\text{train}}/N_{\text{taxa}}$ ratios. **c-d**, The precision-recall (left panels) and receiver operating characteristic (ROC) (right panels) curves for phiNODE applied to synthetic data with the largest sample size generated by the GLV model (top) and DP model (bottom).

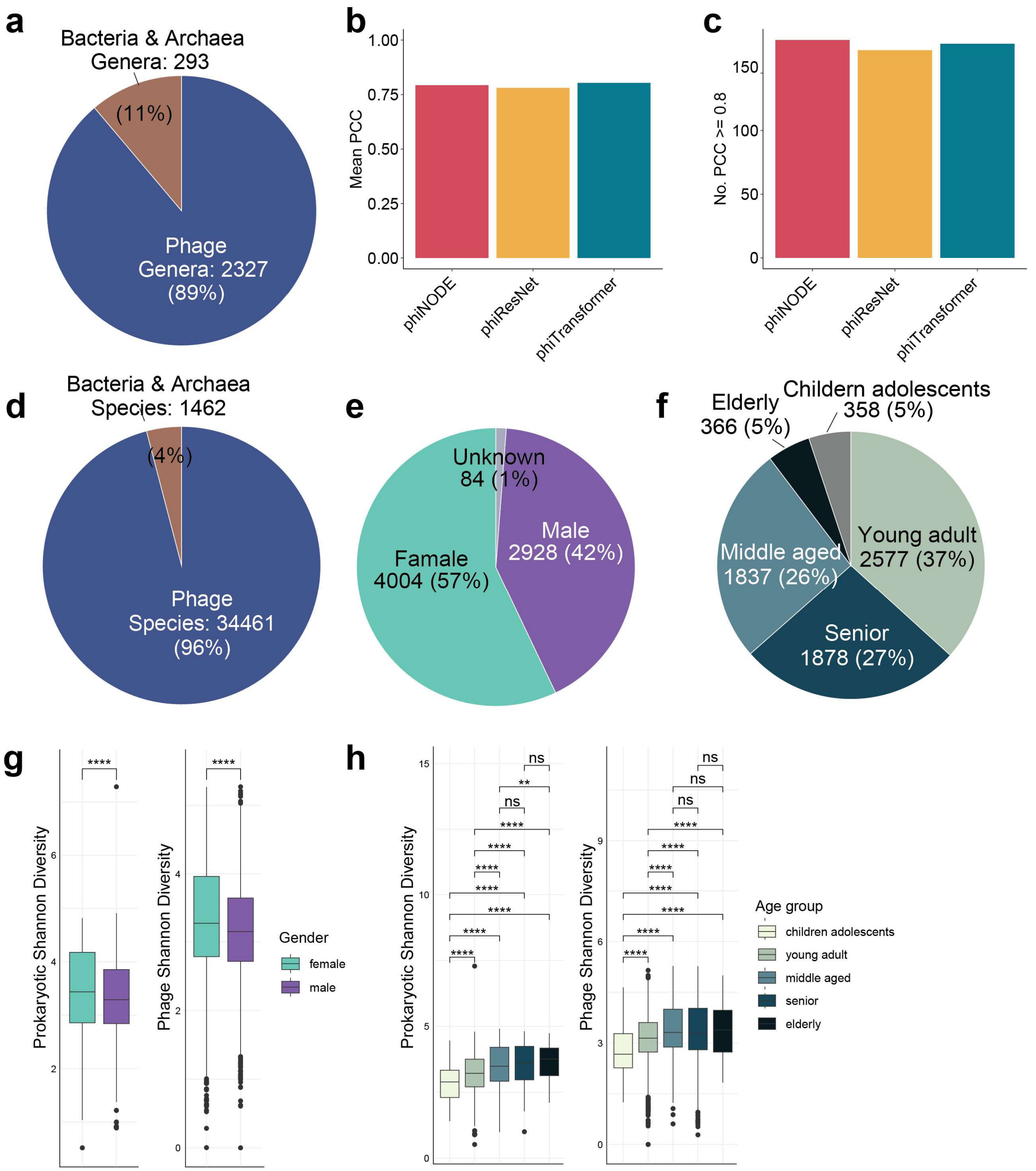


**Supplementary Figure 10**. Training and application of PHILM on 7,016 metagenomic stool samples from healthy individuals. **a**, Distribution of genera detected within the dataset. **b-c**, Performances of phiNODE, phiResNet, and phiTransfromer predicting prokaryotic profiles from phage profiles on test data: (**b**) Mean Pearson correlation coefficient (PCC) values and (**c**) the numbers of prokaryotic genera with PCC values greater than 0.8. **d**, Distribution of species detected within the dataset. **e-f**, Distribution of (**e**) genders and (**f**) age categories within the dataset. **g-h**, Boxplots showing (**g**) prokarytic and (**h**) phage Shannon diversities between different gender and age groups. Significance levels are denoted as follows: ns ($p > 0.01$), ** ($p \leq 0.01$), *** ($p \leq 0.001$), and **** ($p \leq 0.0001$).

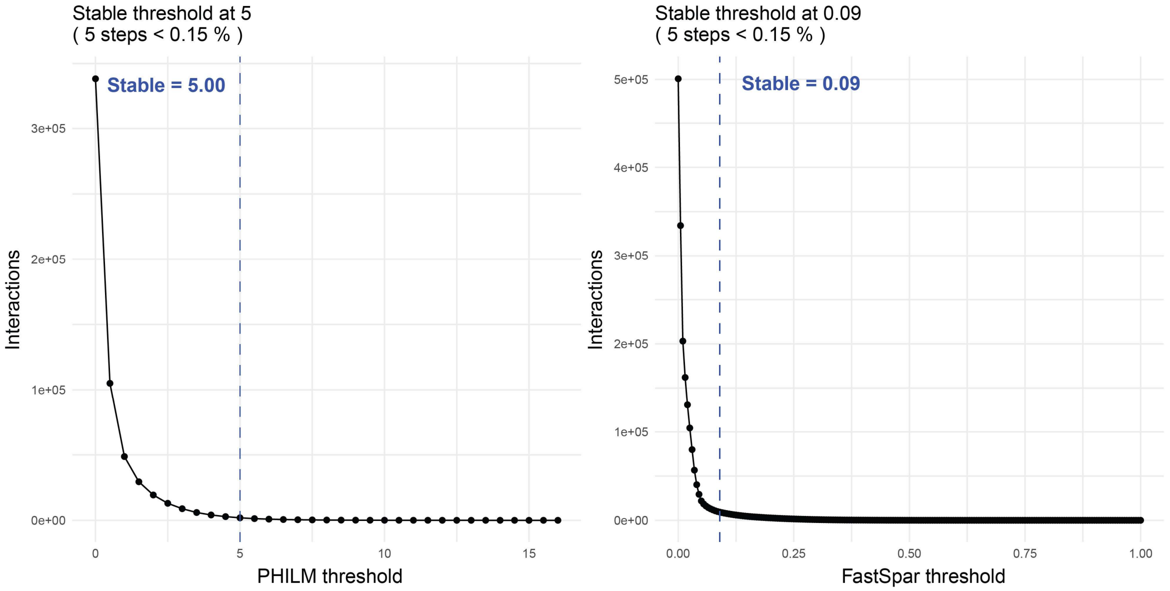


**Supplementary Figure 11**. The line plots depict the number of phage-host interactions recovered at varying sensitivity thresholds for PHILM (using the phiNODE model) (left) and correlation thresholds for FastSpar (right). Vertical dashed lines denote the stabilization thresholds, 5.0 for PHILM and 0.09 for FastSpar, where the relative change in interaction count remains below 0.15% across five consecutive threshold increments.

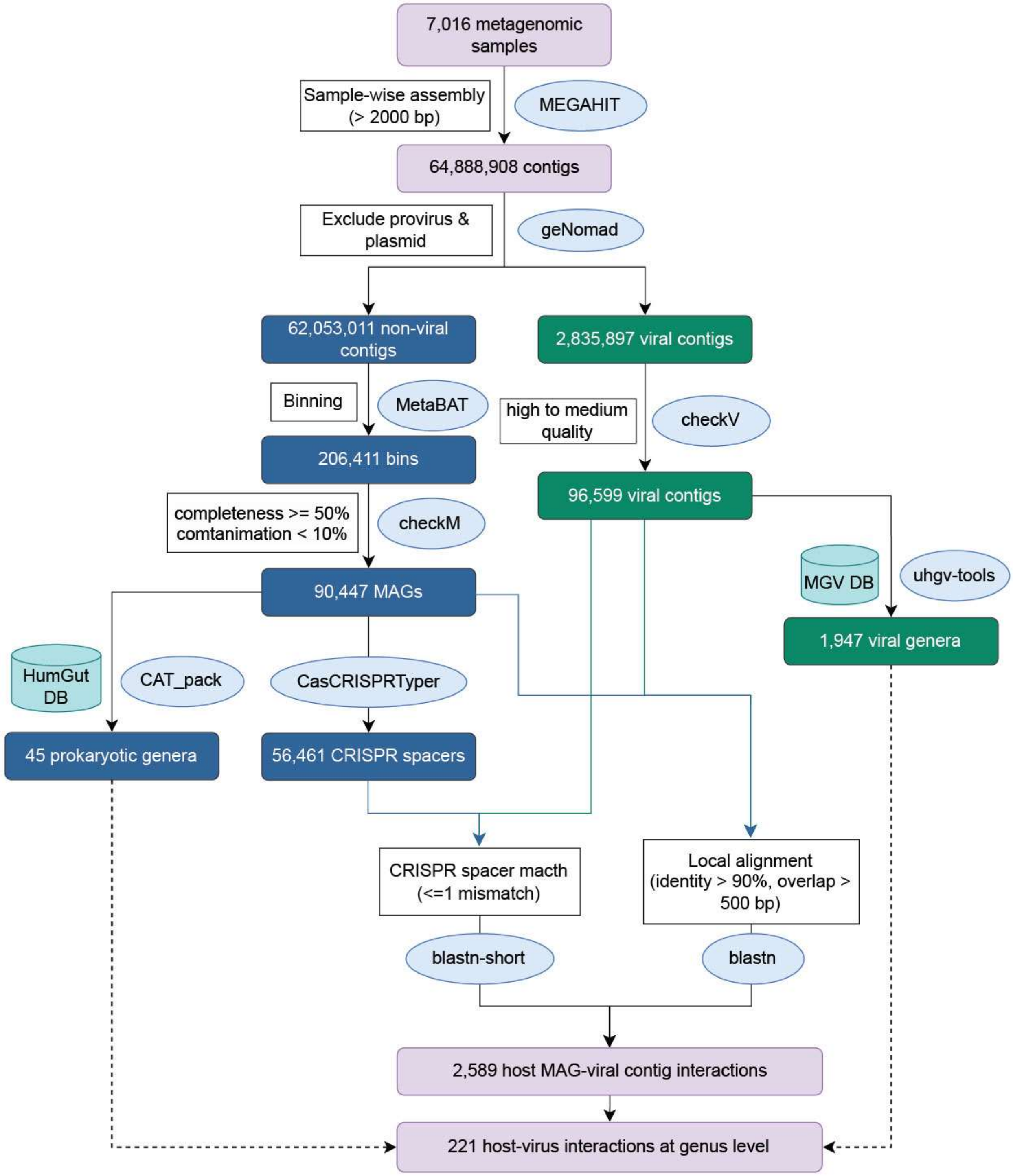


**Supplementary Figure 12**. Workflow for identifying phage-host interactions via contig assembly and mapping of genomic features between prokaryotic metagenome-assembled genomes (MAGs) and viral contigs from 7,016 stool samples of healthy individuals.

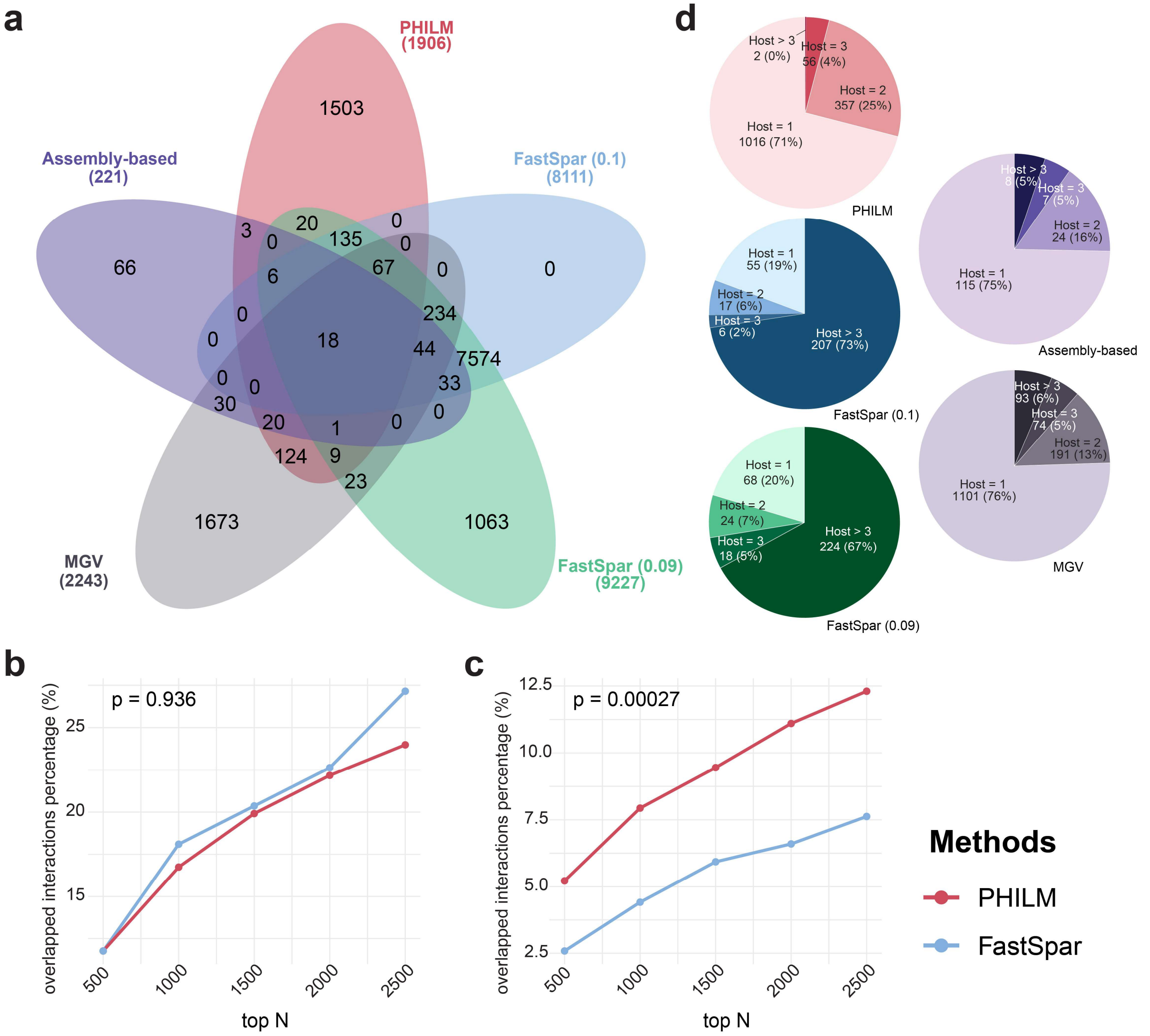


**Supplementary Figure 13**. Overlap of phage-host interactions across multiple methods. **a**, A Venn diagram illustrating the overlaps among phage-host interactions predicted by FastSpar (using ±0.1 and ±0.09 thresholds), the assembly-based method, PHILM, and the interactions recorded in the MGV (Metagenomic Gut Virus) catalogue. **b-c**, Percentages of overlapped genus-level phage-host interactions inferred by PHILM (red) and FastSpar (blue) evaluated at varying top-N cutoffs with those predicted by (**b**) the assembly-based approach and (**c**) the MGV catalogue. P-values were determined by paired t-tests. **d**, Pie charts illustrating the distribution of phage specificities toward the hosts using different approaches.

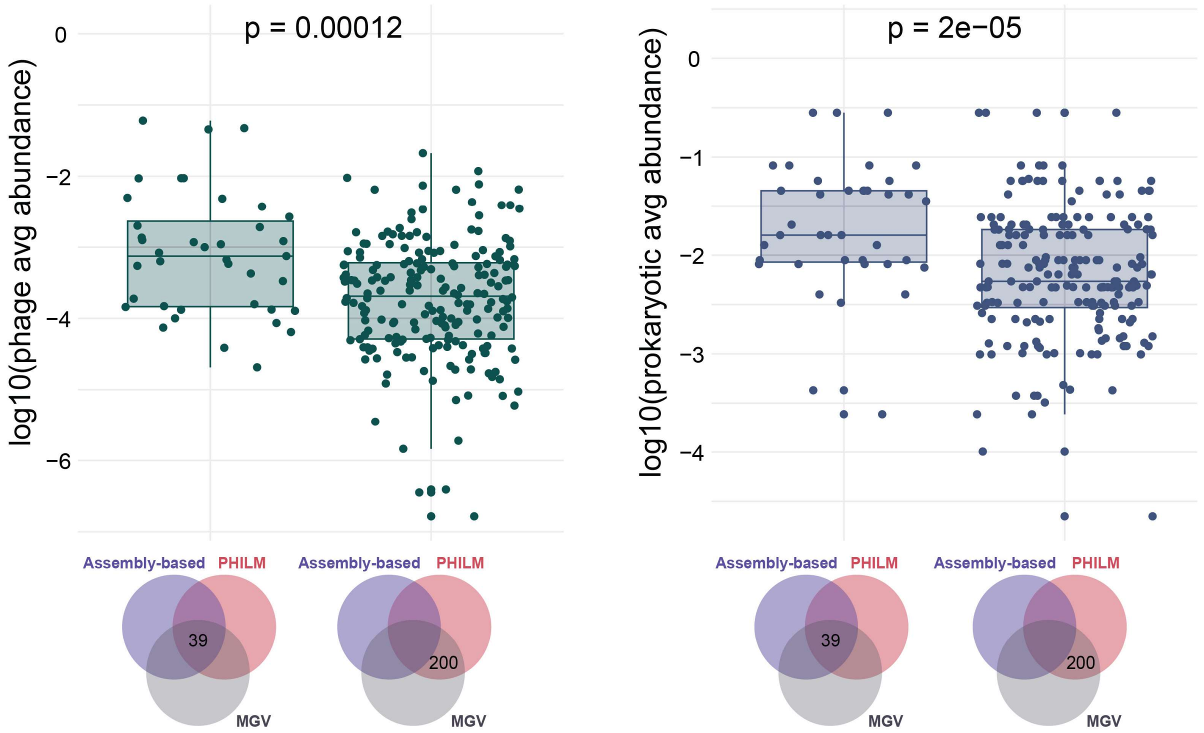


**Supplementary Figure 14**. Boxplots showing the average abundances (with log10 transformation) of phage (left) and prokaryotic (right) genera for interactions shared by PHILM and the MGV catalogue, with (N=39) and without (N=200) overlap with assembly-based interactions. Two-sided Wilcoxon rank-sum tests were used to assess differences between groups.

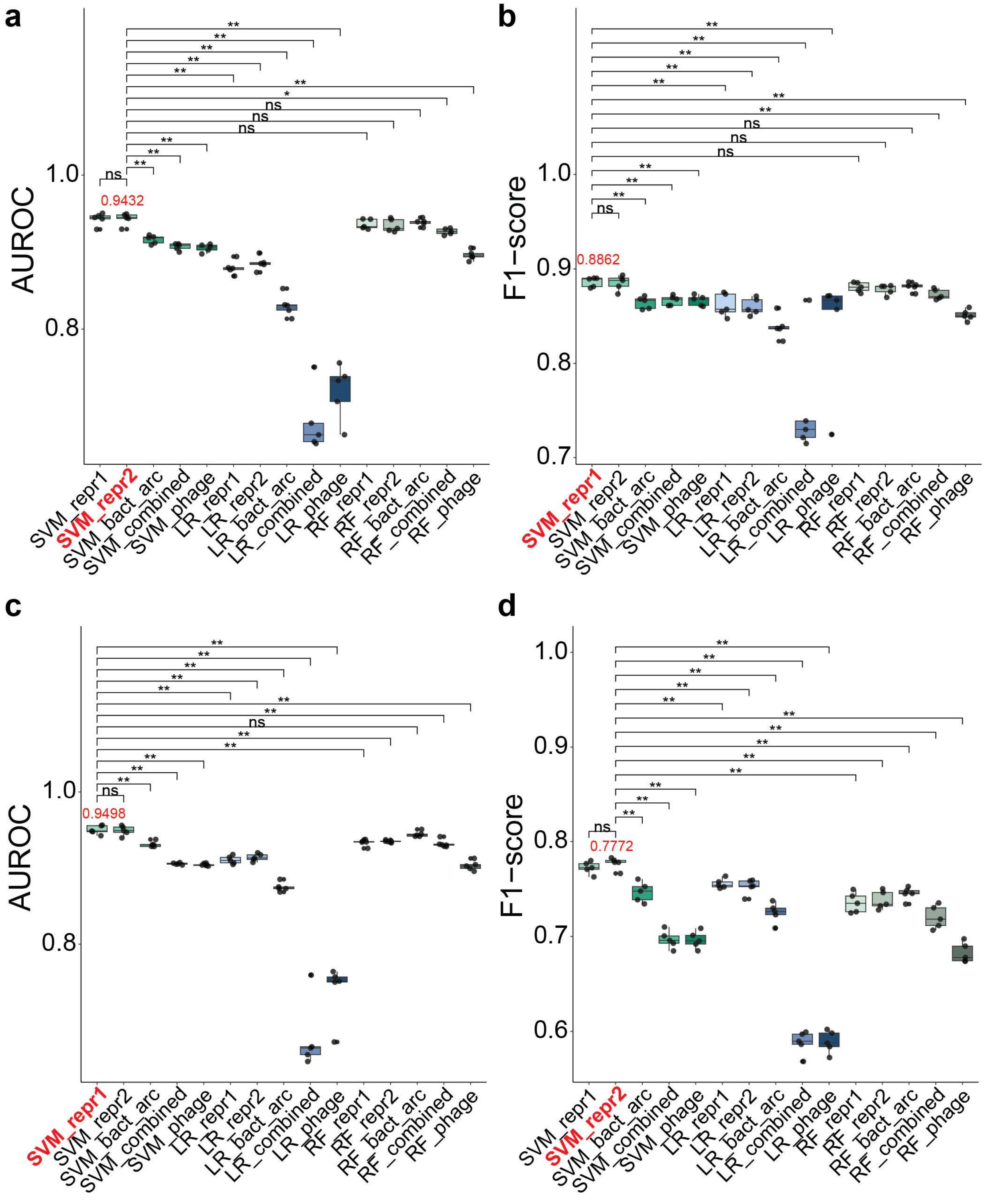


**Supplementary Figure 15**. Performance of all model-feature combination to classify gender and age groups. **a-b**, Boxplots showing the five-cross validation (**a**) AUROCs and (**b**) F1-scores for all model-feature combinations on gender data. **c-d**, Boxplots showing the five-cross validation (**c**) AUROCs and (**d**) F1-scores for all model-feature combinations on age data. The highest values are shown and their corresponding models and features are marked in red. Differences between the best model-feature combination and all other combinations were evaluated using two-sided Wilcoxon rank-sum tests. Significance levels are denoted as follows: ns ($p > 0.05$), * ($p \leq 0.05$) and ** ($p \leq 0.01$).

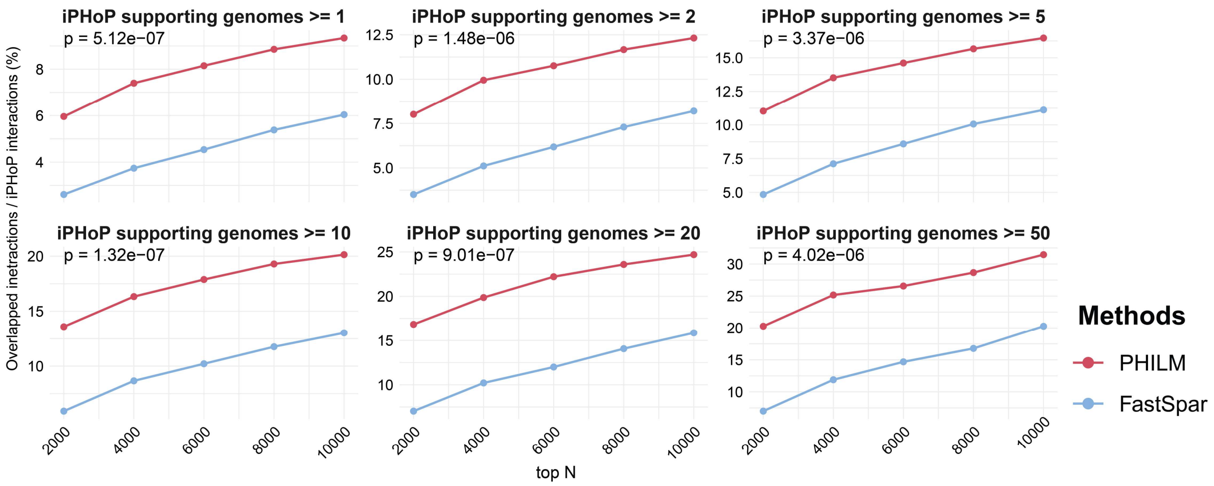


**Supplementary Figure 16**. Percentages of overlapped genus-level phage-host interactions inferred by PHILM (red) and FastSpar (blue) evaluated at varying top-N cutoffs with those predicted by iPHoP and stratified by the number of iPHoP supporting genomes. P-values were determined by paired t-tests.

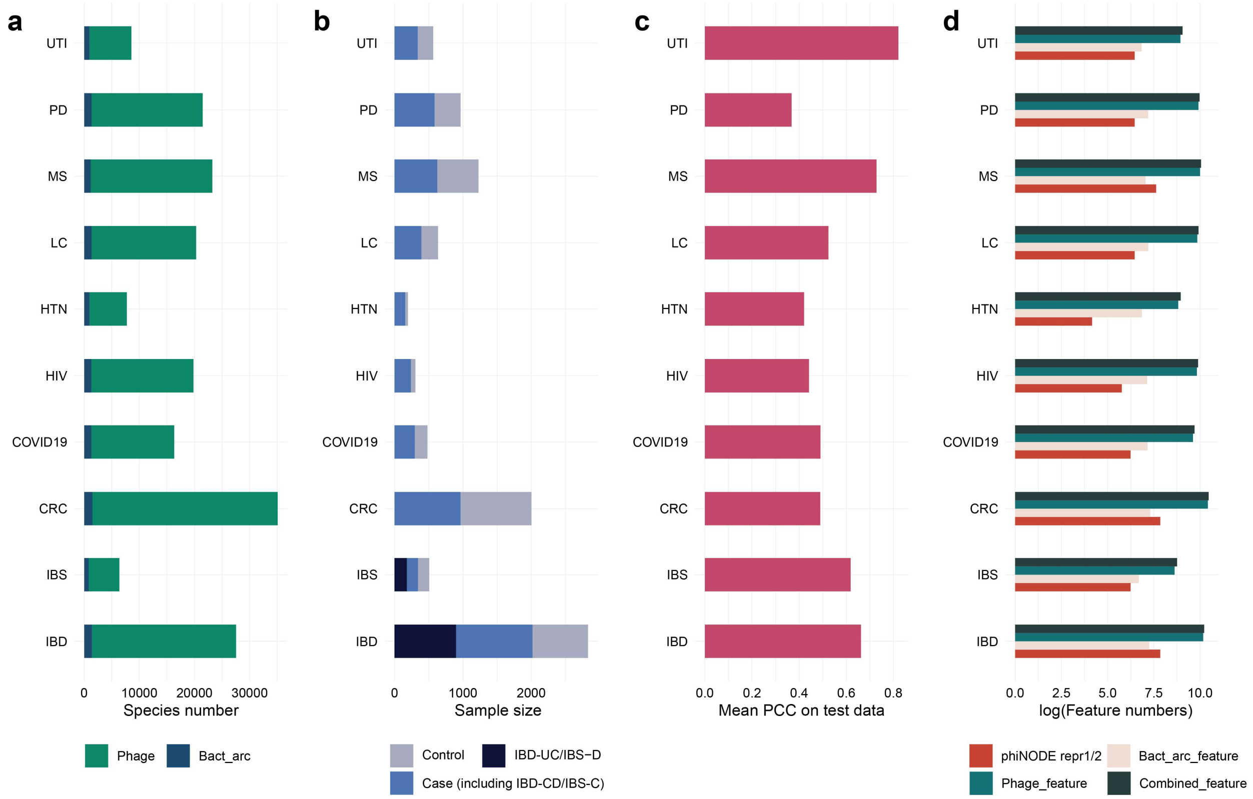


**Supplementary Figure 17**. Basic information for 12 diseases used in disease condition prediction: (**a**) distributions of phage and prokaryotic species, (**b**) distributions of cases and controls, (**c**) mean PCC values, and (**d**) feature numbers for PHILM-derived representations and prokaryotic, phage and combined abudnance-based features.

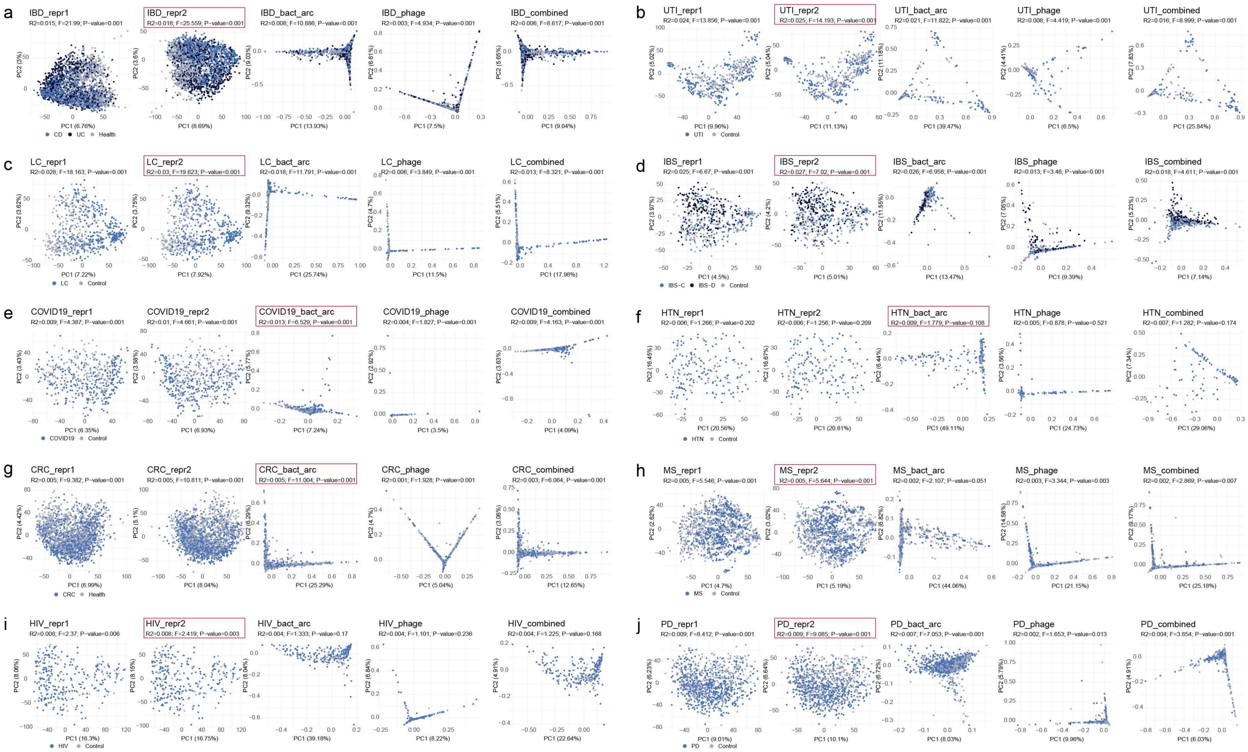


**Supplementary Figure 18**. PCoA plots based on L2 distances for various feature types across 12 diseases (**a-j**). Each plot displays the F values, R² values, and P-values from the PERMANOVA test to quantify differences between case and control groups. Red boxes highlight the features with the highest F values.

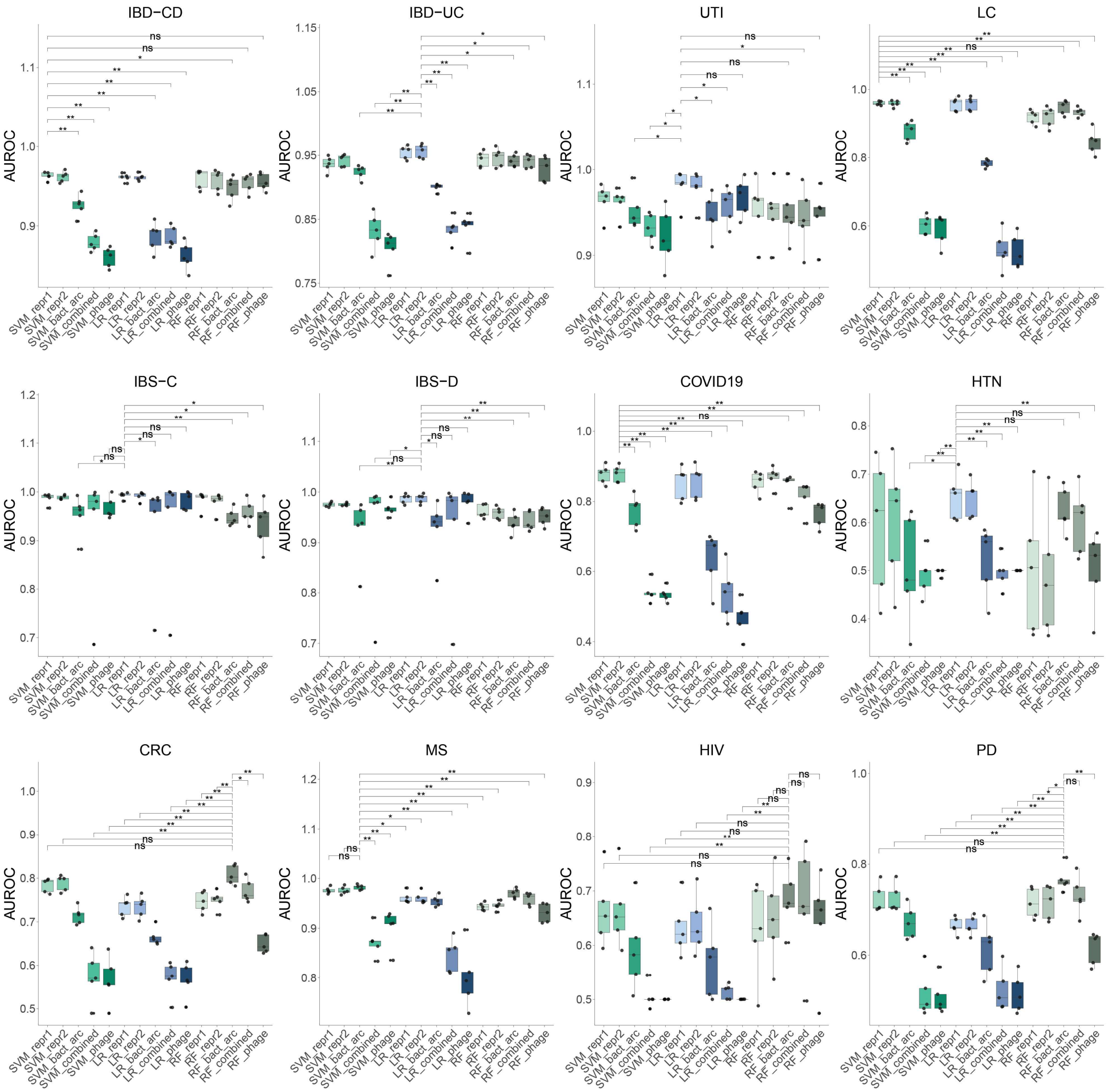


**Supplementary Figure 19**. Boxplots display the five-fold cross-validation AUROCs for all model-feature combinations across 12 diseases. Differences between the best model-feature combination and all other combinations were evaluated using two-sided Wilcoxon rank-sum tests. Significance levels are denoted as follows: ns (p > 0.05), * (p ≤ 0.05) and ** (p ≤ 0.01).

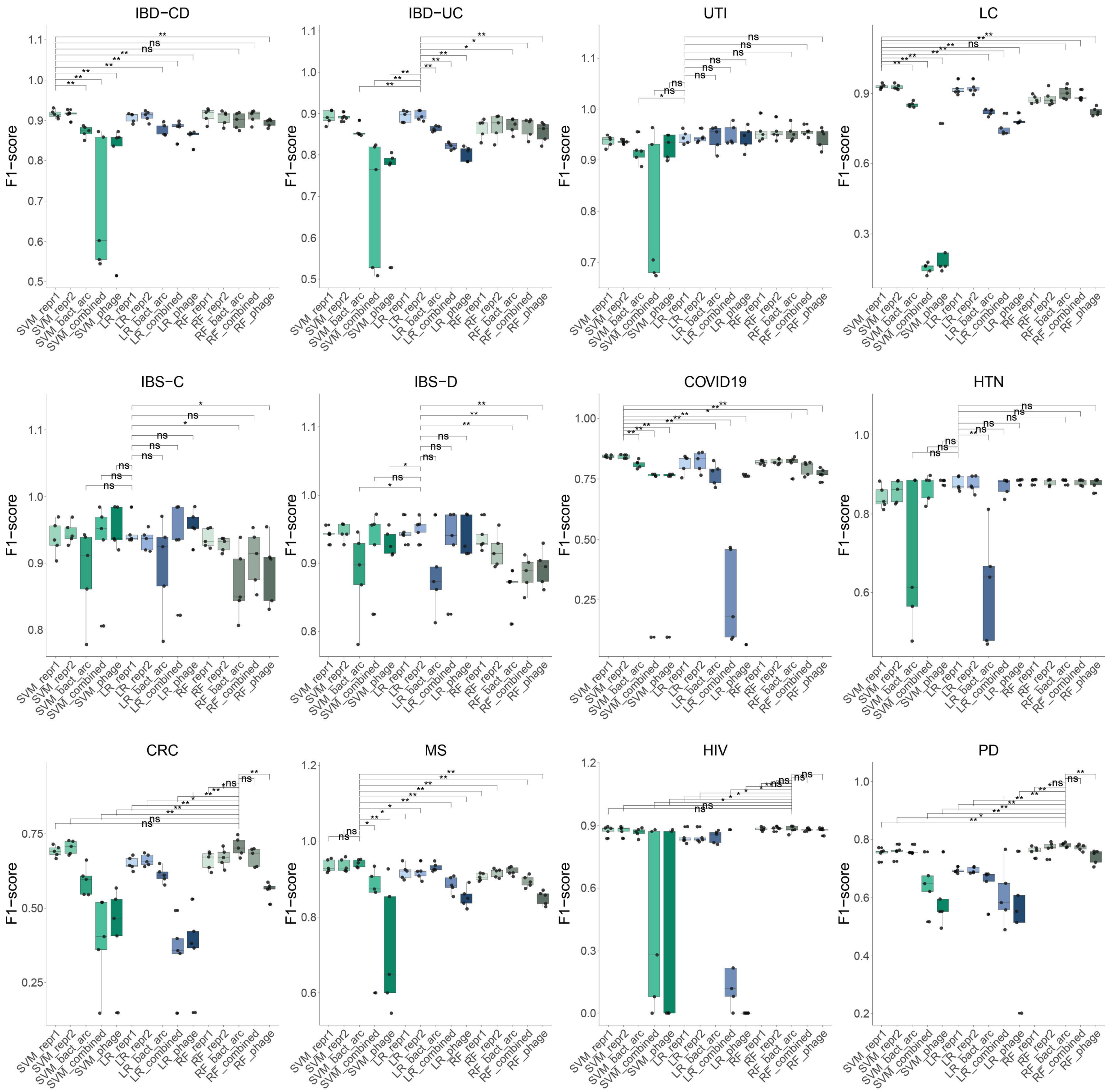


**Supplementary Figure 20**. Boxplots display the five-fold cross-validation F1-scores for all model-feature combinations across 12 diseases. Differences between the best model-feature combination and all other combinations were evaluated using two-sided Wilcoxon rank-sum tests. Significance levels are denoted as follows: ns ($p > 0.05$), * ($p \leq 0.05$) and ** ($p \leq 0.01$).
